# Scarless Enriched selection of Genome edited Human Pluripotent Stem Cells Using Induced Drug Resistance

**DOI:** 10.1101/522383

**Authors:** Keun-Tae Kim, Ju-Chan Park, Haeseung Lee, Hyeon-Ki Jang, Yan Jin, Wankyu Kim, Jeongmi Lee, Hyongbum Henry Kim, Sang-Su Bae, Hyuk-Jin Cha

**Affiliations:** Department of Life Sciences, Sogang University, Seoul 04107, Republic of Korea; College of Pharmacy, Seoul National University, Seoul 08826, Republic of Korea; Ewha Research Center for Systems Biology, Division of Molecular & Life Sciences, Ewha Womans University, Seoul 03760, Republic of Korea; Research Institute for Convergence of Basic Sciences, Hanyang University, Seoul, 04763, Republic of Korea; School of Pharmacy, Sungkyunkwan University, Suwon, Gyeonggi-do 16419, Republic of Korea; Department of Pharmacology, Yonsei University College of Medicine, Seoul, 03722, Republic of Korea; Departement of Chemistry, Hanyang University, Seoul, 04763, Republic of Korea

**Keywords:** Genome editing, CRISPR, Knockout, YM155, *SLC35F2*, Human pluripotent stem cells, drug resistance, Gene editing, CRISPR/Cas9, CCR5, SLC35F2, YM155, human pluripotent stem cells, disease modeling

## Abstract

An efficient gene editing technique for use in human pluripotent stem cells (hPSCs) would have great potential value in regenerative medicine, as well as in drug discovery based on isogenic human disease models. However, the extremely low efficiency of gene editing in hPSCs is a major technical hurdle that remains to be resolved. Previously, we demonstrated that YM155, a survivin inhibitor developed as an anti-cancer drug, induces highly selective cell death in undifferentiated hPSCs. In this study, we demonstrated that the high cytotoxicity of YM155 in hPSCs, which is mediated by selective cellular uptake of the drug, is due to high expression of *SLC35F2* in these cells. Consistent with this, knockout of *SLC35F2* with CRISPR-Cas9 or depletion with siRNAs made hPSCs highly resistant to YM155. Simultaneous gene editing of a gene of interest and transient knockdown of *SLC35F2* following YM155 treatment enabled genome-edited hPSCs to survive because YM155 resistance was temporarily induced, thereby achieving enriched selection of genome-edited clonal populations. This precise and efficient genome editing approach took as little as 3 weeks without cell sorting or introduction of additional genes.

## Introduction

Along with recent advances in human pluripotent stem cells (hPSCs) (Avior et al., 2016; Shi et al., 2017), genome engineering in hPSCs has shown great potential for gene correction in the context of regenerative medicine (Hockemeyer and Jaenisch, 2016). The use of hPSCs derived from patients with genetic disorders enables a constant supply of certain types of cells that mimic the pathology of interest (i.e., disease modeling), and can thus make valuable contributions to both drug screening and basic studies (Engle and Puppala, 2013; Hockemeyer and Jaenisch, 2016). However, patient-derived induced PSCs (iPSCs) require large numbers of case and control iPSCs to minimize the effects of variability among unrelated hPSC lines. By contrast, genome-edited hPSCs provide isogenic pairs of control and disease model cells, enabling rigorous comparisons (Hendriks et al., 2016; Merkle and Eggan, 2013; Musunuru, 2013).

Despite the great potential of genome engineering in hPSCs, the extremely low efficiency of this procedure, along with the time-consuming and laborious procedures required for clonal selection, remains a critical obstacle to a wide range of applications, such as pooled sgRNA screening, which can be performed in mouse ESCs (Koike-Yusa et al., 2014). Another factor contributing to this technical barrier is the fact that Cas9 activity in hPSCs is low (Mali et al., 2013); moreover, the cells undergo massive p53-dependent cell death in response to DNA damage by effect of Cas9 (Ihry et al., 2018). Multiple strategies have been developed to address the low efficiency of genome editing in hPSCs (as well as in other somatic cell models): inducible Cas9 systems (Cao et al., 2016; Gonzalez et al., 2014), puromycin selection (Steyer et al., 2018), self-replicating episomal vectors (Xie et al., 2017), and so on. Alternatively, enriched selection of a minority of the target-edited clones has been attempted by a ‘co-targeting’ approach that enables any target-edited clone to survive through acquired resistance induced by knockout of a gene responsible for drug sensitivity (Agudelo et al., 2017; Liao et al., 2015). However, it would be preferable to achieve ‘scarless enrichment’ of target-edited clones without creating permanent ‘scars’ or introducing foreign genes (e.g., manipulation of a gene responsible for induction of drug resistance), in order to minimize undesirable bias from the additional manipulation.

Teratoma formation resulting from unintended transplantation of undifferentiated hPSCs is a serious risk of hPSC-based cell therapy (Lee et al., 2013a). To address this danger, several approaches have been developed for selective removal of residual hPSCs (Jeong and Cha, 2017). For example, we previously reported that YM155, a survivin inhibitor developed as an anti-cancer drug, is selectively cytotoxic against undifferentiated hPSCs and inhibits teratoma formation (Lee et al., 2013b), and this finding has been reproduced in several independent studies (Bedel et al., 2017; Kang et al., 2017; Kim et al., 2017). At that time, however, the molecular mechanism underlying the high susceptibility of hPSCs to YM155 had not been fully elucidated.

In this study, we found that selective cellular uptake of YM155, which is responsible for the selective cell death of undifferentiated hPSCs, is due to high expression of *SLC35F2*, which encodes a solute carrier membrane transporter. Introduction of gene editing single guide RNA (sgRNA) targeting a gene of interest in parallel with transient siRNA-mediated knockdown of *SLC35F2* efficiently achieved enriched selection of genome-edited hPSCs via induced YM155 resistance. Moreover, this scarless approach to highly efficient enriched selection did not require laborious clonal selection, and thus allowed a single clone to be obtained in as little as 3 weeks.

## Results

### High expression of *SLC35F2* mediates intracellular uptake of YM155 in hPSCs

Selective induction of cell death in hPSCs by YM155, which decreases teratoma formation (Bedel et al., 2017; Kang et al., 2017; Kim et al., 2017; Lee et al., 2013b), is associated with the DNA damage response (Lee et al., 2013b). Cytotoxicity of YM155 cannot be fully prevented by suppression of *BIRC5* (which encodes survivin) (Sim et al., 2017). Hence, we sought to identify the mechanism underlying selective cell death of hPSCs. To this end, we took advantage of bulk cancer cell line gene expression data and drug response data from the Cancer Cell Line Encyclopedia (CCLE) and Cancer Therapeutics Response Portal (CTRP), respectively. First, using data from gene expression omnibus (GEO) (Table. S1), we identified a hPSC signature, i.e., a set of genes that are differentially expressed in hPSCs relative to their differentiated counterparts. Using the hPSC signature, we calculated hPSC scores (Materials and Methods in detail) for 666 human cancer cells by performing single-sample Kolmogorov–Smirnov (KS) enrichment analysis. By correlating the hPSC scores with the sensitivities of each cell line to each of 543 compounds, we identified YM155 as the most effective drug in cells with high hPSC scores (Fig. 1B); this implies that YM155 should be highly specific for hPSCs, as demonstrated previously (Bedel et al., 2017; Kang et al., 2017; Kim et al., 2017; Lee et al., 2013b). Based on correlation of drug and gene expression signature (Fig. 1A), genes whose expression was highly correlated with sensitivity to YM155 were identified as described previously (Rees et al., 2016). *SLC35F2*, a membrane solute carrier, was the transcript most correlated with YM155 sensitivity (Fig. 1B, left panel), suggesting that cellular import of YM155 by *SLC35F2* is responsible for the highly selective cytotoxicity of the drug in hPSCs (Lee et al., 2013b); this is consistent with previous reports in cancer models (Nyquist et al., 2017; Radic-Sarikas et al., 2017; Winter et al., 2014). Using a database of gene expression profiles (http://nextbio.com) (Kupershmidt et al., 2010), we compared relative *SLC35F2* expression between 24 human embryonic stem cells (hESCs) and various cancer cell lines; PC-3, a human prostate cancer cell line, was used as a positive control (P.C.) for high expression of *SLC35F2* (Winter et al., 2014). As shown in Figure 1C, all hESCs (including H9, mainly used in this study) expressed *SLC35F2* at higher levels than the other cancer cell lines, consistent with a previous study that identified hESC surface marker proteins (Kolle et al., 2009). On the other hand, mesenchymal stem cells derived from hESCs (hESC-MSCs) and human dermal fibroblasts (hDFs), which are both highly resistant to YM155 treatment (Kim et al., 2017; Lee et al., 2013b), expressed lower levels of *SLC35F2* than two independent hESC lines (H9 and hCHA3) and induced pluripotent stem cells (iPSCs: SES8) (Lee et al., 2013b) (Figs. 1D and E). SLC35F2 is the membrane transporter responsible for uptake of YM155 into cells, resulting in DNA damage (Winter et al., 2014); consistent with this, marked DNA damage was detected solely in hESCs, but not in hESC-MSCs, following YM155 treatment (Figs. 1F and G). The higher intracellular level of YM155 in hESCs relative to hDFs (Fig. 1H) clearly indicates that high *SLC35F2* expression in hPSCs results in selective cytotoxicity of this drug in hPSCs.

**Figure 1.**
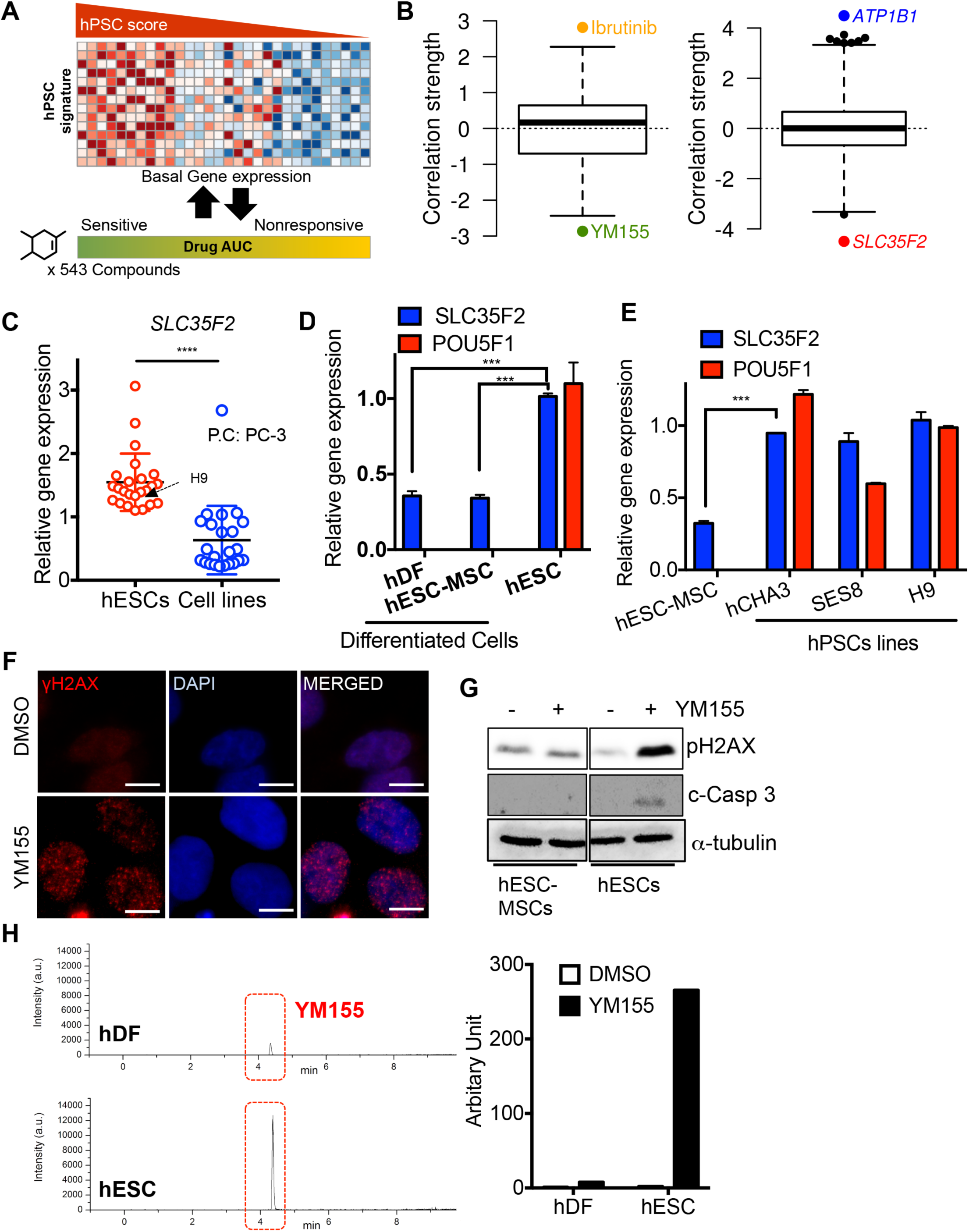
High expression of *SLC35F2* mediates intracellular uptake of YM155 in hPSCs. (A) A graphical schema of correlating cell line hPSC scores with cell line drug responses (AUC) against each of 543 compounds available in CTRP (B) Left, the correlation strength (z-scored Pearson correlation coefficient) of 543 compounds from comparing AUCs of each compound to hPSC scores, Cells with high hPSC scores (hPSC-like cells) are selectively sensitive to YM155. Right, the correlation strength of 18,858 genes from comparing the expression values of each gene with AUCs of YM155. *ATP1B1* and *SLC35F2* are highly expressed in YM155 resistant and sensitive cells, respectively. (C) Expression of *SLC35F2* in several hESC cell lines from NextBio portal, Differentiated cells (blue dots) were analyzed as controls. (P.C. indicates PC-3 prostate cancer cell). (D) mRNA expression of *SLC35F2* and *POU5F1* of human dermal fibroblasts (hDF), hESC-derived MSCs and hESCs (E) mRNA expression levels of *SLC35F2* and *POU5F1* of hESC-MSCs, hCHA3, H9 and SES8 iPSCs. (F) Fluorescent images of H9 stained with γH2AX after YM155 treatment, DAPI for nuclear counterstaining (scale bars = 10 µm) (G) Immunoblotting assay between hESC-MSCs and hESCs were performed to detect pH2AX (Ser 139) and cleaved Caspase 3 (c-Casp 3) level after YM155 treatment. α-tubulin was used for internal control. (H) LC-MS/MS analysis was performed to quantify the intracellular YM155 amount between hDF and hESCs. 1 µM of YM155 was treated for 2 hours.

### SLC35F2 is responsible for YM155-induced selective cell death in hPSCs

To explore the connection between the selective cytotoxicity of YM155 in hPSCs (EC_50_: 10 nM, Fig. S1B) and the high expression of *SLC35F2* in these cells (Figs. 1D, E, and S1A), we knocked out *SLC35F2* in hESCs using CRISPR/Cas9 by targeting exon 7, as previously described (Winter et al., 2014) (Fig. 2A). After introducing a single guide RNA (sgRNA) against *SLC35F2* along with Cas9, we treated the cells with YM155 to ablate wild-type hESCs that had not been edited. Following YM155 treatment, a few colonies survived, whereas most cells underwent cell death (Fig. 2B, red arrow). One of the surviving colonies (Fig. 2B, YM155-resistant clone: YM155R) was selected and maintained. Clone YM155R was highly resistant to further YM155 treatment (Fig. 2B, right panel). Furthermore, after introduction of the sgRNA against *SLC35F2* and Cas9, insertion/deletion (indel) frequency was increased in a concentration-dependent manner (Fig. 2C), and in the presence of 100 nM YM155, the surviving clones were nearly 100% genome-edited hESCs (100% indel) (Fig. 2D), suggesting that the *SLC35F2* knockout (*SLC35F2* KO) hESC population was enriched in a dose-dependent manner, thereby efficiently eliminating false positives (i.e., hESCs that survived with wild-type *SLC35F2*). A single clone from YM155R maintained under YM155 treatment turned out to be a homozygous bi-allelic *SLC35F2* KO (Fig. 2E). This clone (*SLC35F2* KO hESCs: KO#1) was highly resistant to YM155-induced cell death (Fig. 2F and Movie S1). However, like control hESCs, it underwent dramatic cell death in response to bleomycin (BLM), which is imported by SLC22A16, another member of the solute carrier family (Aouida et al., 2010) (Fig. S1C). We concluded that the acquired resistance of *SLC35F2* KO hESCs to YM155 was the result of low intracellular levels of YM155 due to failure of drug uptake (Fig. 2H) and subsequent fewer DNA damage response in the absence of *SLC35F2* (Fig. 2G).

**Figure 2.**
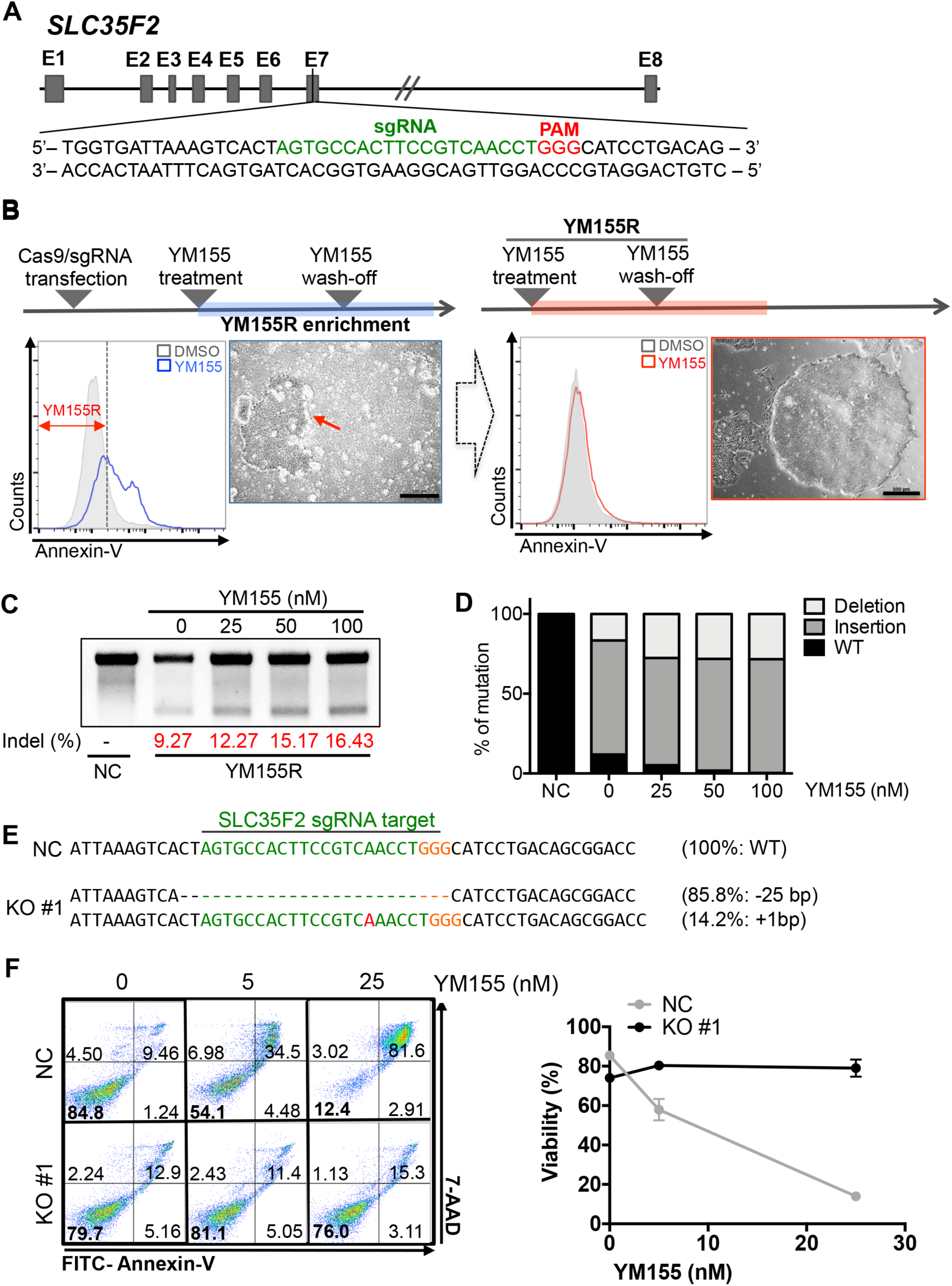
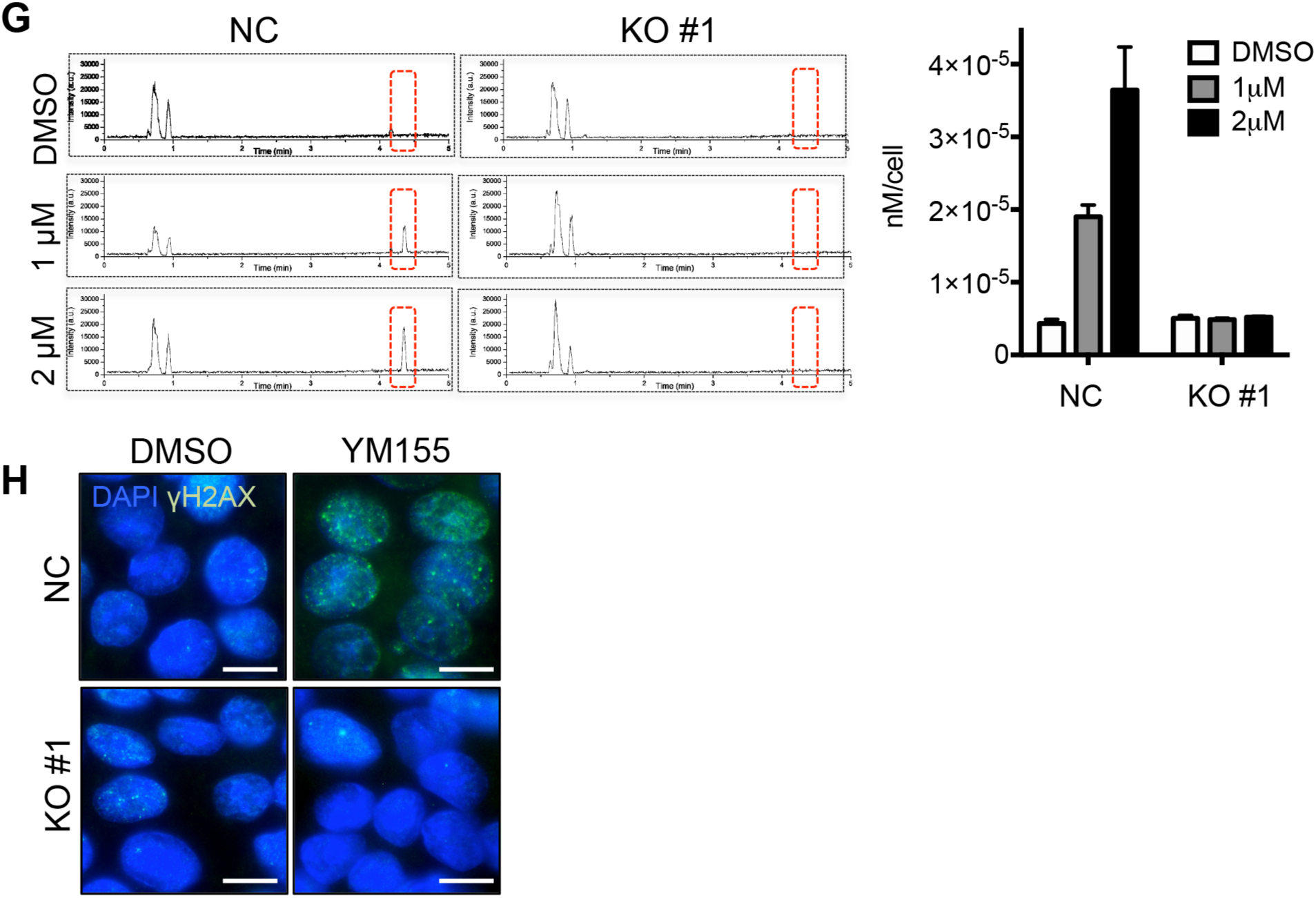
*SLC35F2* enables YM155-induced selective cell death in hPSCs. (A) Scheme for SLC35F2 knockout targeting Exon 7 (B) Experimental schema of enrichment of SLC35F2 KO hESCs (top panel), Flow cytometry of Annexin V and microscopic images of hESCs after 50nM of YM155 treatment (Red arrow indicates YM155 resistant clone: YM155R, scale bars = 500 µm) (C) T7E1 assay with sgRNA of SLC35F2 and Cas9 transfected hESCs after treatment of indicative dose of YM155, Indel percentage shown as below (D) Percentage of mutation quantified from NGS analysis was graphically presented. (E) Sequence information from a YM155R clone (KO #1), sgRNA target sequences and PAM sequence were shown in Red and Orange respectively. (F) Flow cytometry of Annexin-V apoptosis assay in WT (NC) and SLC35F2 KO#1 hESCs after indicative dose of YM155 treatment (G) Intracellular level of YM155 in control (NC) and KO hESCs (KO#1) determined by LC-MS/MS (H) Fluorescent microscopic images of hESCs stained with γH2AX in control (NC) and SLC35F2 KO (KO#1) hESCs with DMSO as a vehicle or 20 nM of YM155 treatment, DAPI for nuclear counterstaining (scale bars = 10 µm)

To characterize the *SLC35F2* KO hESCs, we monitored the expression levels of typical pluripotency markers such as *NANOG*, *SOX2*, and *POU5F1*. Consistent with the absence of alteration of typical pluripotency markers (Fig. 3A), protein levels of SOX2, OCT4, and LIN28A (Figs. 3B and S1D), as well as alkaline phosphatase activity (Fig. 3C), were not significantly altered in *SLC35F2* KO hESCs. Consistent with this, self-renewal of *SLC35F2* KO hESCs was also maintained, as reflected by their growth rate (Figs. 3D and S1E), fluorescence-based competition assay with WT control (Fig. 3E) (Cha et al., 2017), and *in vitro* and *in vivo* differentiation potentials (Figs. 2F and G). In support of these findings, comparison of transcriptome data from WT and KO hESCs revealed a linear relationship in the expression levels of pluripotency genes (Korkola et al., 2006) (red circles) (Fig. 3H).

**Figure 3.**
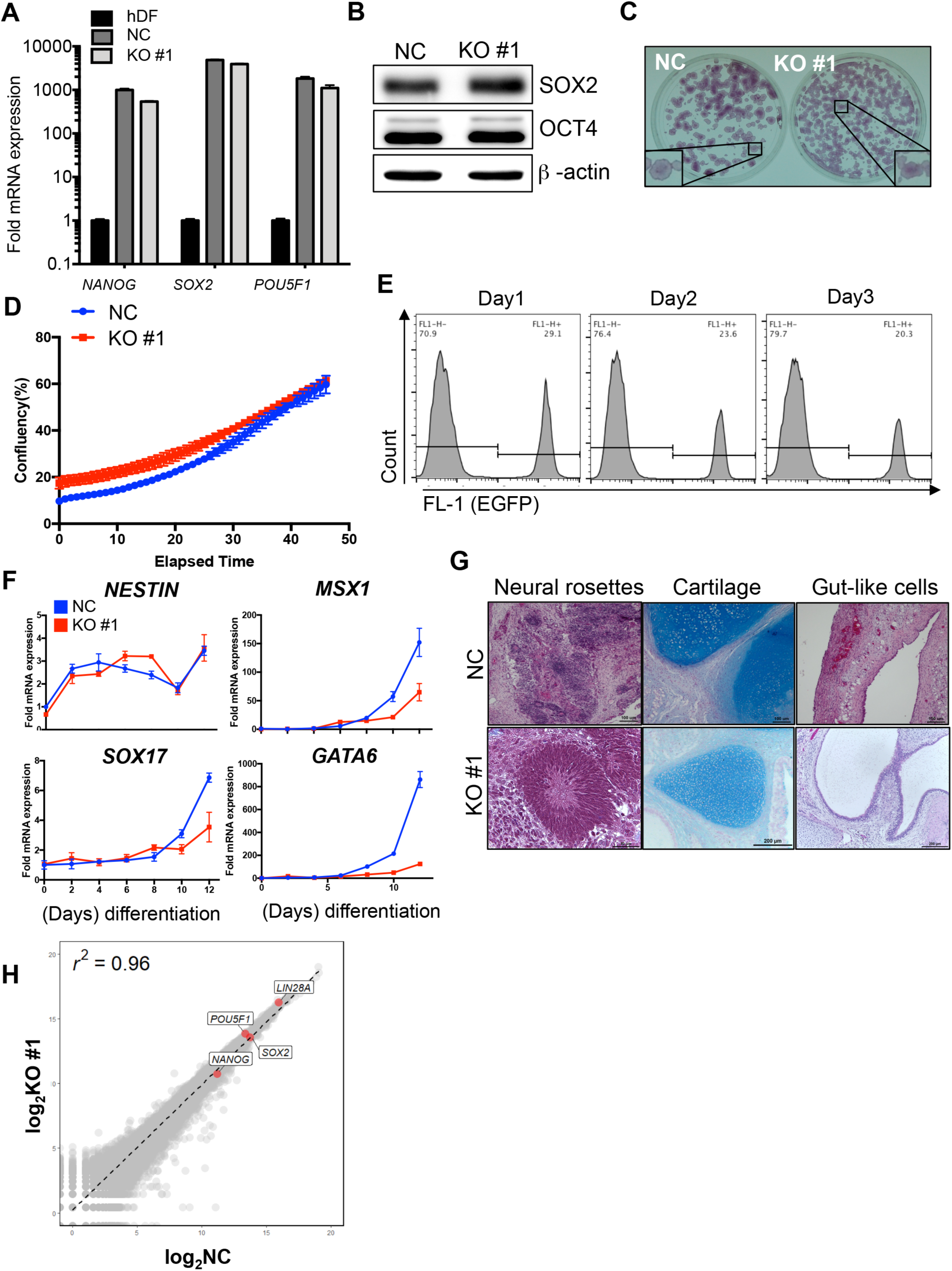
*SLC35F2* KO hPSCs maintain pluripotency and proliferation intact. (A) Relative mRNA expression of indicative pluripotent markers in human dermal fibroblast, and control (NC) and SLC35F2 KO (KO#1) hESCs (B) Immunoblotting analysis for indicative proteins (SOX2 and OCT4 in control (NC) and SLC35F2 KO (KO#1) hESCs, β–actin was used as a loading control. (C) Alkaline phosphatase activity assay of control (NC) and SLC35F2 KO (KO#1) hESCs, Insets are enlarged images of representative colonies, respectively. (D) Cell proliferation rate of control (NC) and SLC35F2 KO (KO#1) hESCs was graphically presented. (E) Flow cytometry of green fluorescence of EGFP expressing control and SLC35F2 KO (KO#1) hESCs determined at indicative days (F) Relative levels of mRNA expression of typical lineage specific genes (endoderm: *SOX17, GATA6*, mesoderm: *MSX1* and ectoderm: *NESTIN*) of control (NC) and SLC35F2 KO (KO#1) hESCs at indicative days after *in vitro* differentiation (G) Images of typical three germ layers from the teratoma stained with H&E staining, Masson’s trichrome, and Alcian Blue. The scale bar represents 50 mm. (H) Scatter plot of whole genes from control (NC) and SLC35F2 KO (KO#1) hESCs, Typical pluripotent marker genes (*POU5F1*, *SOX2*, *NANOG*, *Lin28A*) were marked as red circles.

### YM155-mediated enriched selection

The observation that enriched selection of *SLC35F2* KO hESCs could be achieved by YM155 treatment led us to predict that induced resistance to YM155 could be useful for isolating genome-edited hPSCs if *SLC35F2* were co-targeted with the gene of interest (GOI) (Fig. 4A). For a proof-of-concept study, we took advantage of HEK293T cells, which express relatively high levels of *SLC35F2* (Fig. S2A), leading to dose-dependent cell death following YM155 treatment (10-fold higher EC_50_ than for hESCs, Fig. S2B). Simple introduction of sgRNA for *SLC35F2* along with Cas9, followed by YM155 treatment, led to establishment of a genome-edited population (Fig. 4B) at up to 88.4% efficiency (Fig. 4C). Like hESCs, *SLC35F2* KO HEK293T cells were highly resistant to YM155, but not doxorubicin (Fig. S2C and D), due to failure of cellular uptake of YM155 (Fig. S2E).

**Figure 4.**
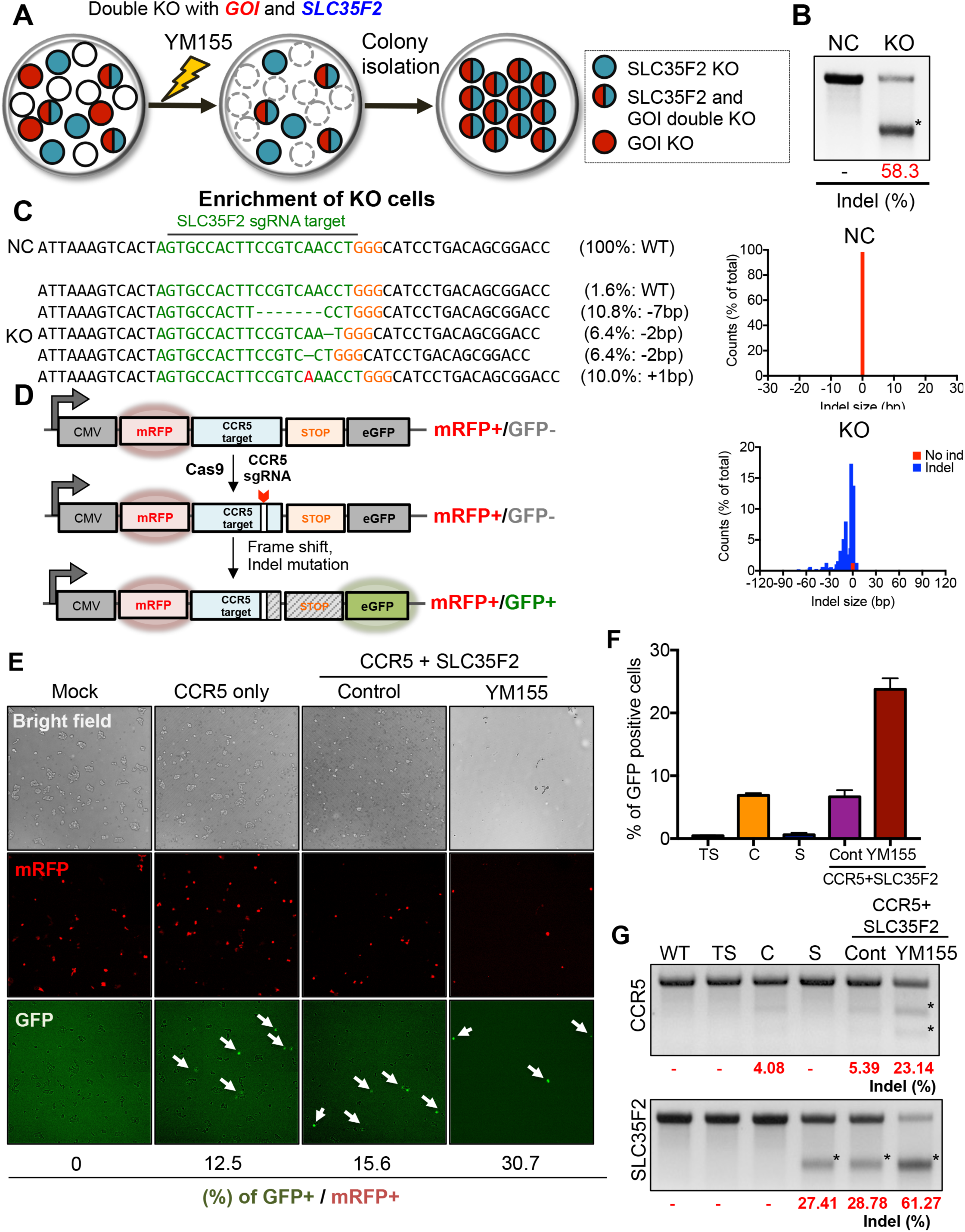
YM155 mediated enriched selection. (A) Graphical schema for YM155 mediated enriched selection approach in HEK293T cells (GOI: Gene-of-interest) (B) T7E1 assay of control (NC) and SLC35F2 KO HEK293T cells, The asterisk indicates a cleaved band after T7E1 enzyme treatment. (C) NGS data of control (NC) and SLC35F2 KO (KO#1) HEK293T cells, Indel variety of control (NC) and SLC35F2 KO (KO) HEK293T cells was graphically presented. (D) A graphical summary of surrogate vector system for CCR5 target (E) Fluorescence microscopic images of HEK293T cells expressing surrogate reporter after sgRNA transfection of *CCR5* only (CCR5 only) or *CCR5/SLC35F2* (CCR5+SLC35F2) with or without YM155 treatment, Ratio of GFP over mRFP positive population was shown in below. Average of ratio of GFP over mRFP positive population at indicative conditions (TS: Surrogate reporter only, C: sgRNA for CCR5, S: sgRNA for SLC35F2, CCR5+SLC35F2: sgRNA for both CCR5 and SLC35F2) was graphically presented. (G) T7E1 assay for CCR5 and SLC35F2 at indicative conditions (TS: Surrogate reporter only, C: sgRNA for CCR5, S: sgRNA for SLC35F2, CCR5+SLC35F2: sgRNA for both CCR5 and SLC35F2), The asterisk indicates a cleaved band after T7E1 enzyme treatment. Relative band densities were shown in below.

Next, we attempted co-targeting of *SLC35F2* along with the *CCR5*. In these experiments, we took advantage of the *CCR5* surrogate reporter system (Ramakrishna et al., 2014) to monitor the efficiency of *CCR5* targeting. In this system, the monomeric red fluorescent protein (mRFP) is expressed constitutively, whereas expression of green fluorescent protein (GFP) is initiated only after *CCR5* is targeted by Cas9 expression (Fig. 4D). Thus, the proportion of GFP-positive cells enables live monitoring of CRISPR/Cas9 targeting efficiency. After co-targeting *SLC35F2* and *CCR5*, followed by YM155 treatment, we observed a significant increase in the GFP-positive population (Figs. 4E and F). Considering that the surrogate reporter system reveals only one-third of the genome-edited population (Ramakrishna et al., 2014), the ~25% GFP-positive population following YM155 selection implied a genome-edited population as high as 80% (Fig. 4F). It is also noteworthy that YM155 treatment eliminated most mRFP-expressing cells (red box), thereby selectively enriching for GFP-positive cells (blue box) (Fig. S2F, Movies S1 and S2). To confirm enrichment of *CCR5* KO by co-targeting of *SLC35F2* followed by YM155 treatment, we performed the T7 endonuclease 1 (T7E1) assay for *CCR5*; the results revealed a distinct increase in KO efficiency (Fig. 4G).

### YM155-based enriched selection of genome-edited hESCs

After testing this approach (YM155-based Enriched Selection of CRISPR Co-target; hereafter, ‘YES-approach’) in HEK293T cells (Fig. 4), we applied it to hPSCs, which are more sensitive to YM155 (Figs. 1 and 2). For proof-of-concept of the YES-approach, we attempted to produce *CCR5*-targeted hESCs. We chose this gene for the following reasons: (i) disruption of *CCR5* (a genomic safe harbor (Sadelain et al., 2011)) has little effect on the pluripotency of hPSCs (Kang et al., 2015); and (ii) the cells would have potential future applications in HIV-1 research, e.g., production of *CCR5*-depleted CD4^+^ T cells (Holt et al., 2010; Hutter et al., 2009; Lai, 2012). In these experiments, we used two different ratios of sgRNAs targeting *CCR5* and *SLC35F2* to minimize the selection of false-positive clones (e.g., *CCR5*WT/*SLC35F2*KO) that could still survive under YM155 treatment. Through the YES-approach with sgRNA targeting *CCR5* (C) and *SLC35F2* (S), seven (for C:S=1:1 ratio) or four (C:S=2:1) colonies survived and were isolated. Among them, three clones and one clone, respectively, were *CCR5*-targeted hESCs (over 25% efficiency) (Fig. 5A). Accordingly, a single clone after the selection was shown as 85.5% indel occurred, implying that the surviving clones were successfully genome-edited in the desired manner (Fig. 5B).

**Figure 5.**
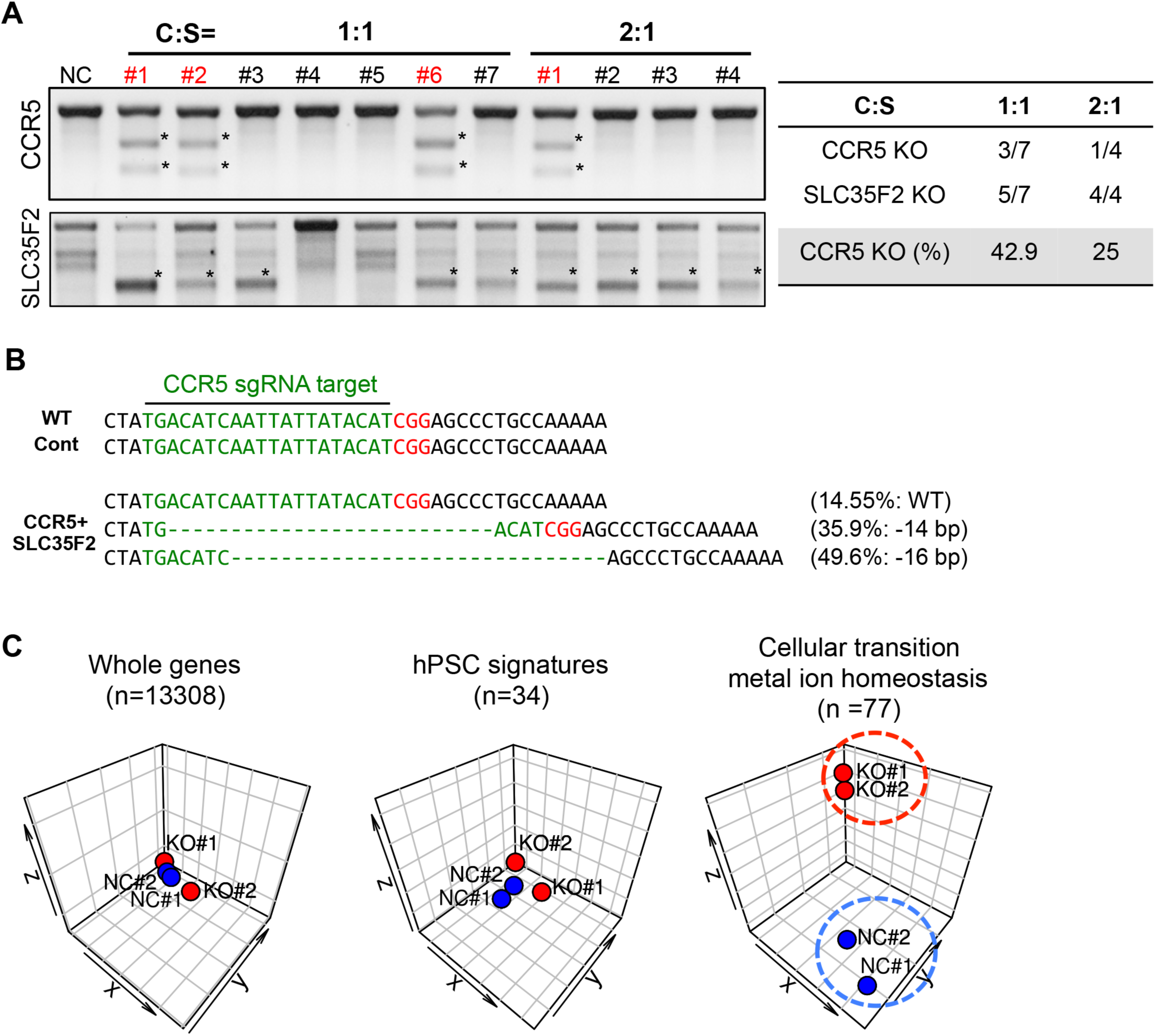
YM155 based enriched selection of genome-edited hESCs. (A) T7E1 assay for CCR5 and SLC35F2 in each clone of hESCs (Seven clones for C:S=1:1, and four clones for C:S=2:1) at indicative conditions {different dose ratio of sgRNA of CCR5(C) vs SLC35F2(S)} (left panel), Summary table of number of KO clones of CCR5 and SLC35F2 (right panel) (B) NGS result from #1 clone (from C:S=1:1) compared to control (WT control), sgRNA target and PAM sequence were shown in Green and Red letters respectively. (C) Clustering of RNA-seq samples using t-distributed stochastic neighbor embedding (t-SNE) based on the expression of whole genes, hPSC signature genes, and cellular transition metal ion homeostasis (GO:0046916) genes.

Although overall gene expression and expression of pluripotency genes were not altered in *SLC35F2* KO hESCs (Fig. 3H), we could not rule out the effect of permanent KO of *SLC35F2* in hESCs. Therefore, we closely examined the genes whose expression was altered in *SLC35F2* KO cells. Although we detected only minor differences between wild-type (NC#1 and NC#2) and KO cells (KO#1 and KO#2) in both overall transcription and the hPSC signature, expression of genes involved in ‘cellular transition metal ion homeostasis,’ which are associated with the original function of *SLC35F2* as a membrane transporter, was altered significantly (Fig. 5C and Table. S2).

### Scarless YES-approach for establishment of *CCR5*-targeted hESCs

Although the YES-approach was effective for establishing GOI-targeted hESCs, permanent KO of *SLC35F2* to induce YM155 resistance would be problematic from the standpoint of isogenic disease modeling because the effects of *SLC35F2* on mineral or ion homeostasis (Fig. 5C and Table S2) could not be ruled out. To address this complication, transient induced YM155 resistance mediated by depletion of *SLC35F2* was achieved by introduction of a siRNA against *SLC35F2* along with the sgRNA targeting the GOI (in this case, *CCR5*) (Fig. 6A). As predicted, depletion of *SLC35F2* had fully recovered 5 days after siRNA transfection (Fig. 6B). Induced YM155 resistance disappeared at the same time (Fig. 6C, right panel), allowing for enriched selection at 2 days after siRNA transfection (Fig. 6C, left panel). These observations suggest that induced YM155 resistance mediated by siRNA against *SLC35F2* occurred transiently, with no permanent scar.

**Figure 6.**
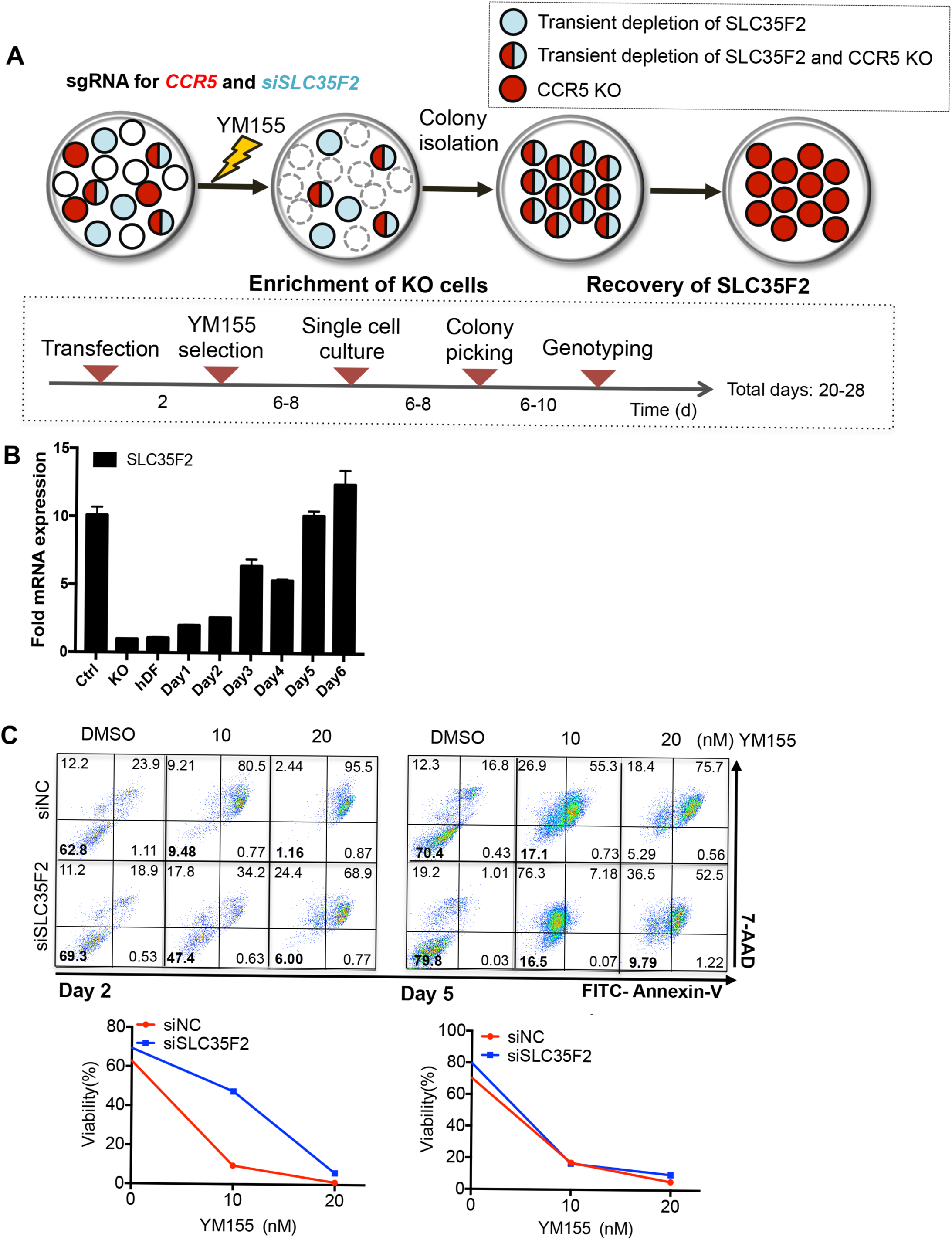
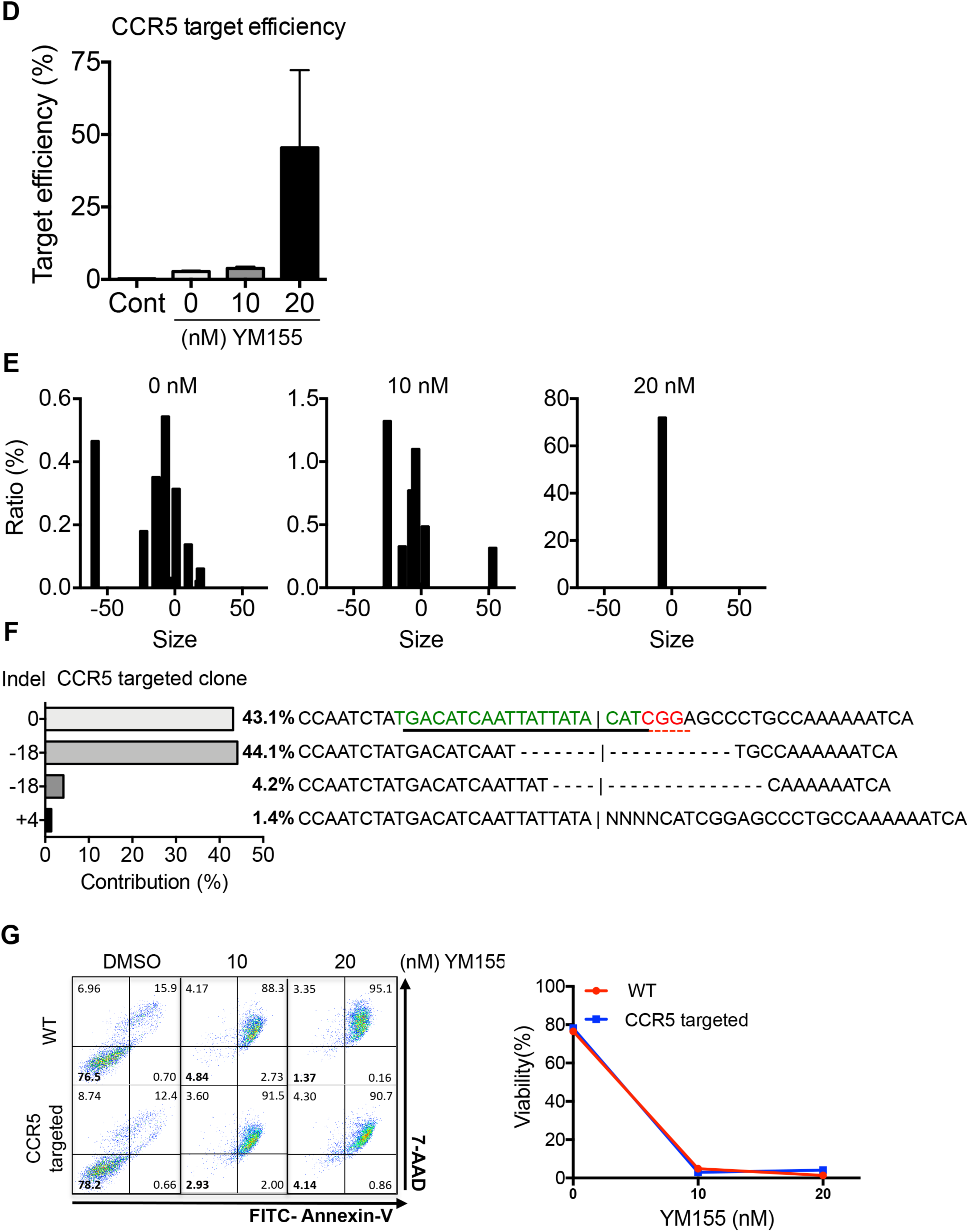
Scarless YES-approach for establishment of *CCR5* targeted hESCs. (A) Graphical schema of scarless YES-approach (B) Relative levels of mRNA expression of *SLC35F2* at indicative days after siRNA transfection in hESCs (C) Flow cytometry of Annexin V and 7AAD staining at 2 or 5 days after transfection of control siRNA (siNC) and siRNA for SLC35F2 (siSLC35F2) with indicative dose of YM155 treatment (D) Average of ratio of indel in CCR5 after YES-approach at indicative dose of YM155 treatment (E) Deep sequencing analysis of CCR5 KO clones achieved by YES-approach at indicative dose of YM155 treatment (F) NGS data for CCR5 target sequence, sgRNA target and PAM sequence were shown in Green and Red respectively. (G) Flow cytometry of Annexin-V/7-AAD staining at indicative dose of YM155 in WT or CCR5 KO hESCs

We then applied this scarless YES-approach to *CCR5* gene editing in hESCs. As shown in Figure 6D, *CCR5* target efficiency was improved in a YM155 dose-dependent manner. Notably in this regard, the indel percentage in samples with the YES-approach reached 72.2% even without laborious isolation of numerous single colonies (Figs. S3A-D). Accordingly, the mutation patterns of the genome-edited population were simplified by treatment with 20 nM YM155, indicating that few colonies survived the treatment (Figs. 6E and S3E). One of the clones obtained from the YES-approach was correctly targeted to CCR5 (Figs. 6F and S3F). Also, the CCR5 targeted clone maintained pluripotency as determined by alkaline phosphatase activity and expression level of pluripotency markers (Fig. S3G). Along with recovery of *SLC35F2* expression (Fig. S3G), the cytotoxicity of YM155 was fully restored (Figs. 6G and S3H), suggesting the functional recovery of *SLC35F2*.

## Discussion

In order to avoid the laborious procedures made necessary by the extremely low efficiency of genome editing using current techniques, a variety of strategies have been developed to improve editing efficiency not only in hPSCs (Gonzalez et al., 2014; Mitzelfelt et al., 2017; Steyer et al., 2018), but also in somatic cell models (Agudelo et al., 2017; Cao et al., 2016; Liao et al., 2015). Among them, enriched selection approaches based on inducing drug resistance by targeting the causative gene responsible for drug sensitivity (Agudelo et al., 2017; Liao et al., 2015) would be advantageous because selection of desirable clones results from altered phenotype (e.g., induced drug resistance) caused by effect of gene editing unlike puromycin selection, corresponding to merely Cas9 expression (Steyer et al., 2018). In these procedures, however, induced drug resistance was achieved by producing a permanent scar in the causative genes (Agudelo et al., 2017; Liao et al., 2015), potentially resulting in unexpected bias in hPSCs-based disease models.

Here, we demonstrated that selective induction of cell death in hPSCs by YM155 (Lee et al., 2013b) results from selective cellular uptake of the drug due to high expression of *SLC35F2* (Figs. 1 and S1A). The high cytotoxicity of YM155 was completely abolished by knockout of *SLC35F2* (Fig. 2), and the induction of YM155 resistance by knockout of *SLC35F2* was useful for enriched selection of genome-edited clones (Figs. 4 and 5). Live monitoring revealed that un-targeted cells underwent cell death in response to YM155 (Fig. S2F, Movies S2 and S3). Considering the undesirable biases that could result from permanent knockout of *SLC35F2* (Fig. 5), we then attempted to temporarily induce YM155 resistance by transient knockdown of *SLC35F2*, and we found that this was sufficient to achieve enriched selection of targeted clones (Fig. 6). The function of *SLC35F2* in the edited clones was fully recovered (Fig. 6). Following this scarless YES-approach, CRISPR-targeted clones were achieved with up to 72.2% efficiency, avoiding the need for laborious clonal selection within 3 weeks.

It is important to note that YM155 sensitivity varies depending on cell type due to differences in the expression level of *SLC35F2* (Fig. S2A) as well as the sensitivity of the DNA damage response. The high expression level of *SLC35F2* in most hPSCs (Kolle et al., 2009) (Fig. 1C) may render the YES-approach uniquely suitable for application in these cells. In addition, the strong mitochondria-dependent apoptotic cell death response of hPSCs by genotoxic stresses (Liu et al., 2013), which depends on mitochondrial translocation of p53 (Lee et al., 2013b), may contribute to the high selectivity of YM155: as previously reported (Winter et al., 2014), intracellular YM155 taken up through SLC35F2 induced a clear DNA damage response (Fig. 1G). Unlike other chemical-based selection approaches, induced YM155 resistance was achieved by abrogating a drug-specific membrane importer, thereby blocking cellular uptake of YM155. Thus, due to their failure to take up YM155, clones that survived enriched selection would be free from undesirable effects of YM155.

However, the scarless YES-approach requires colony selection to avoid false-positive clones (Fig. 5A) that could survive YM155 treatment. Notwithstanding, development of a one-vector system for expression of siRNA and Cas9 would minimize the incidence of false-positive or -negative clones. Taken together, our findings indicate that the scarless YES-approach could be further developed as a useful tool for advancing the application of genome engineering in hPSCs. In particular, this strategy would enable the use of sgRNA libraries to establish whole panels of hPSCs that mimic genetic diseases.

## Acknowledgments

This work was supported by a grant from the National Research Foundation of Korea (NRF-2017R1A2A2A05000766 2017M3A9B3061843 and 2017M3C9A5028691).

## Author Contributions

HJ.C conceived the overall study design and led the experiments. KT.K and JC.P mainly conducted the experiments, data analysis, and critical discussion of the results. H.L and W.K conducted the informatics analysis. HK.J and H.K and SS.B contributed to sgRNA design and NGS analysis. Y.J and J.L measured the intracellular level of YM155. All authors contributed to manuscript writing and revising, and endorsed the final manuscript.

## Declaration of Interests

The authors declare no competing interests.

## STAR Methods

### Pharmacogenomics data sets

Basal-level cell line mRNA expression data was obtained from the Cancer Cell Line Encyclopedia (CCLE) data portal (https://portals.broadinstitute.org/ccle/data) (Barretina et al., 2012). Raw Affymetrix CEL files were converted to normalize real expression value for each gene probe set using Robust Multi-array Average (RMA) method. When collapsing probes to genes, the maximum of expression values of multiple probes per gene was taken as a representative expression value for the gene. Among the total 886 cell lines, we excluded hematologic lineages, non-human, and ambiguous tissue types, and used 666 cancer cell lines with drug sensitivity data in this analysis. Cell line drug response data was downloaded from Cancer Therapeutics Response Portal (https://ocg.cancer.gov/programs/ctd2/data-portal)(Seashore-Ludlow et al., 2015). Cell viability values were adjusted to a range of 0-100 % and fitted by 4-parameter logistic regression analysis. We selected a unique range of concentrations that tested on the most cells for each compound to calculate AUC (area under the fitted curve). AUC was normalized to a range of 0-1 by the maximum AUC assumed to be 0 % growth inhibition in a given concentration range.

### Cell line and culture

Human embryonic stem cells (WA09, WiCell Research Institute) were cultured on Matrigel (BD Biosciences) coated dishes fed with mTeSRTM-E8TM media (STEMCELL technologies) and StemMACSTM media (Miltenyi-Biotec) added with 50 µg/ml Gentamicin (Life Technologies). Cells were passaged every 5-6 days and media was changed every day. hESC were rinsed with DPBS and exposed to Dispase (Gibco) for detachment. Detached cells were washed with DMEM/F-12 (Gibco) media and plated on Matrigel-coated plate. 10 µM of Y27632 (Gibco) was added for cell attachment when needed. HEK293T cells were cultured on a culture dish (Falcon) fed with DMEM (Gibco) added with 10% FBS (Gibco) and 50 µg/ml Gentamicin. Upon transfer, HEK293T cells were rinsed with DPBS and detached enzymatically with 0.25% Trypsin. Trypsin was inactivated by adding DMEM containing 10% FBS and appropriate cells were seeded on the dishes.

### Teratoma formation assay

After anesthesia, direct testicular injection of WT, *SLC35F2* KO hESCs was performed to examine in vivo pluripotency of each cell. 5 × 106 cells of each WT (n=3) and *SLC35F2* KO (n=3) hESC were inoculated subcutaneously and intracutaneously to mice with 4-weeks-old. Mice were euthanized 2 months after inoculation of each WT and *SLC35F2* KO hESC. These animal experiments were conducted under the permission of Seoul National University Institutional Animal Care and Use Committee (permission number: SNU-180810-1).

### RNA-sequencing analysis

Total RNA was extracted from cells using Easy-BLUE^TM^ RNA isolation kit (iNtRON Biotechnology) and quality control was performed. The qualified samples proceed to library construction. The sequencing library is prepared by random fragmentation of the DNA or cDNA sample, followed by 5” and 3” adapter ligation. The read quality of raw RNA-seq files (FASTQ) was checked using FastQC (v0.11.7) and the adapter sequences in reads were eliminated using cutadapt (v1.8.1). Trimmed FASTQ files were aligned to GRCh38 genome using STAR aligner (v2.6.0a), then the read counts for genes were acquired using HTseq (v0.11.0) with Gencode v22 annotation GTF (https://www.gencodegenes.org/human/release_22.html).

### LC-MS/MS analysis

HEK293T and H9 cells were exposed to YM155 for 1 hour, cells were 1 × 106 cells were counted. Harvested cells were lysed with 80% methanol and incubated in ice for an additional 1 hour. After spin down with 13,000 rpm for 20 minutes, supernatants were collected and evaporated with N2 gas until no more solvents remained. Sample residues were re-suspended with 100 µL of 50% methanol followed by 10 seconds sonication, 5 seconds vortex, spin-down and filtration with 0.2 µm membrane filter. Total 5 µL of solvents were injected to UHPLC/Q-TOF MS (Waters, Milford, MA, USA) using ACQUITY UPLC BEH C18 column (50 × 2.1 mm, 1.7 mm, 30 °C). Samples were read with 0.2 mL/ min flow rate and the estimated peak for YM155 was 4.32 ~ 4.9 min.

### T7E1 assay

PCR is performed using SolgTM Taq DNA Polymerase (STD16-R500, SolGent), following the supplier’s instruction with 10 µl of total volume per sample. 1st PCR is performed with primer F1 and primer R for each gene. The 1st PCR product is diluted in 190 µl of DW. 2nd PCR is performed with 1 µl of diluted 1st PCR product, using primer F2 and primer R for each gene. The 2nd PCR product is mixed with the same volume of 2X NEBuffer2 (B7002S, New England BioLabs) and hybridized. 3 units of T7E1 endonuclease (M0302S, New England BioLabs) were treated to 10 µl of hybridized 2nd PCR product. The enzyme reaction is conducted for 40 minutes at 37 °C water bath.

### Flow cytometry

Cells were detached with AccutaseTM (561527, BD Biosciences) and washed with DPBS for three times and then stained with FITC Annexin-V (556419, BD Biosciences) and 7-AAD (559925, BD Biosciences) for cell death detection. Diluted 1X Annexin V binding buffer (556454, BD Biosciences) is used as the solvent for staining. FACScalibur from BD Biosciences and CellQuest software was used for FACS analysis.

### Statistical analysis

The quantitative data are expressed as the mean values ± standard error of the mean (SEM). Student’s paired t-tests or one-way ANOVAs were performed to analyze the statistical significance of each response variable. Pre-specified comparisons between groups were conducted (when appropriate) by Tukey’s post hoc test using the SPSS program (Statistical Package for the Social Sciences, version 17). p values less than 0.05 were considered statistically significant.

## References

Agudelo, D., Duringer, A., Bozoyan, L., Huard, C.C., Carter, S., Loehr, J., Synodinou, D., Drouin, M., Salsman, J., Dellaire, G., et al. (2017). Marker-free coselection for CRISPR-driven genome editing in human cells. Nat Methods 14, 615–620.

Aouida, M., Poulin, R., and Ramotar, D. (2010). The human carnitine transporter SLC22A16 mediates high affinity uptake of the anticancer polyamine analogue bleomycin-A5. J Biol Chem 285, 6275–6284.

Avior, Y., Sagi, I., and Benvenisty, N. (2016). Pluripotent stem cells in disease modelling and drug discovery. Nat Rev Mol Cell Biol 17, 170–182.

Barretina, J., Caponigro, G., Stransky, N., Venkatesan, K., Margolin, A.A., Kim, S., Wilson, C.J., Lehar, J., Kryukov, G.V., Sonkin, D., et al. (2012). The Cancer Cell Line Encyclopedia enables predictive modelling of anticancer drug sensitivity. Nature 483, 603–607.

Bedel, A., Beliveau, F., Lamrissi-Garcia, I., Rousseau, B., Moranvillier, I., Rucheton, B., Guyonnet-Duperat, V., Cardinaud, B., de Verneuil, H., Moreau-Gaudry, F., et al. (2017). Preventing Pluripotent Cell Teratoma in Regenerative Medicine Applied to Hematology Disorders. Stem cells translational medicine 6, 382–393.

Cao, J., Wu, L., Zhang, S.M., Lu, M., Cheung, W.K., Cai, W., Gale, M., Xu, Q., and Yan, Q. (2016). An easy and efficient inducible CRISPR/Cas9 platform with improved specificity for multiple gene targeting. Nucleic acids research 44, e149.

Cha, Y., Han, M.J., Cha, H.J., Zoldan, J., Burkart, A., Jung, J.H., Jang, Y., Kim, C.H., Jeong, H.C., Kim, B.G., et al. (2017). Metabolic control of primed human pluripotent stem cell fate and function by the miR-200c-SIRT2 axis. Nat Cell Biol 19, 445–456.

Engle, S.J., and Puppala, D. (2013). Integrating human pluripotent stem cells into drug development. Cell stem cell 12, 669–677.

Gonzalez, F., Zhu, Z., Shi, Z.D., Lelli, K., Verma, N., Li, Q.V., and Huangfu, D. (2014). An iCRISPR platform for rapid, multiplexable, and inducible genome editing in human pluripotent stem cells. Cell stem cell 15, 215–226.

Hendriks, W.T., Warren, C.R., and Cowan, C.A. (2016). Genome Editing in Human Pluripotent Stem Cells: Approaches, Pitfalls, and Solutions. Cell stem cell 18, 53–65.

Hockemeyer, D., and Jaenisch, R. (2016). Induced Pluripotent Stem Cells Meet Genome Editing. Cell stem cell 18, 573–586.

Holt, N., Wang, J., Kim, K., Friedman, G., Wang, X., Taupin, V., Crooks, G.M., Kohn, D.B., Gregory, P.D., Holmes, M.C., et al. (2010). Human hematopoietic stem/progenitor cells modified by zinc-finger nucleases targeted to CCR5 control HIV-1 in vivo. Nature biotechnology 28, 839–847.

Hutter, G., Nowak, D., Mossner, M., Ganepola, S., Mussig, A., Allers, K., Schneider, T., Hofmann, J., Kucherer, C., Blau, O., et al. (2009). Long-term control of HIV by CCR5 Delta32/Delta32 stem-cell transplantation. N Engl J Med 360, 692–698.

Ihry, R.J., Worringer, K.A., Salick, M.R., Frias, E., Ho, D., Theriault, K., Kommineni, S., Chen, J., Sondey, M., Ye, C., et al. (2018). p53 inhibits CRISPR-Cas9 engineering in human pluripotent stem cells. Nat Med 24, 939–946.

Jeong, H.C., and Cha, H.J. (2017). Helping Induced hPSCs Clean Up Their Act. Cell chemical biology 24, 651–652.

Kang, H., Minder, P., Park, M.A., Mesquitta, W.T., Torbett, B.E., and Slukvin, II (2015). CCR5 Disruption in Induced Pluripotent Stem Cells Using CRISPR/Cas9 Provides Selective Resistance of Immune Cells to CCR5-tropic HIV-1 Virus. Mol Ther Nucleic Acids 4, e268.

Kang, S.J., Park, Y.I., Hwang, S.R., Yi, H., Tham, N., Ku, H.O., Song, J.Y., and Kang, H.G. (2017). Hepatic population derived from human pluripotent stem cells is effectively increased by selective removal of undifferentiated stem cells using YM155. Stem cell research & therapy 8, 78.

Kim, K.T., Jeong, H.C., Kim, C.Y., Kim, E.Y., Heo, S.H., Cho, S.J., Hong, K.S., and Cha, H.J. (2017). Intact wound repair activity of human mesenchymal stem cells after YM155 mediated selective ablation of undifferentiated human embryonic stem cells. Journal of dermatological science 86, 123–131.

Koike-Yusa, H., Li, Y., Tan, E.P., Velasco-Herrera Mdel, C., and Yusa, K. (2014). Genome-wide recessive genetic screening in mammalian cells with a lentiviral CRISPR-guide RNA library. Nature biotechnology 32, 267–273.

Kolle, G., Ho, M., Zhou, Q., Chy, H.S., Krishnan, K., Cloonan, N., Bertoncello, I., Laslett, A.L., and Grimmond, S.M. (2009). Identification of human embryonic stem cell surface markers by combined membrane-polysome translation state array analysis and immunotranscriptional profiling. Stem Cells 27, 2446–2456.

Korkola, J.E., Houldsworth, J., Chadalavada, R.S., Olshen, A.B., Dobrzynski, D., Reuter, V.E., Bosl, G.J., and Chaganti, R.S. (2006). Down-regulation of stem cell genes, including those in a 200-kb gene cluster at 12p13.31, is associated with in vivo differentiation of human male germ cell tumors. Cancer Res 66, 820–827.

Kupershmidt, I., Su, Q.J., Grewal, A., Sundaresh, S., Halperin, I., Flynn, J., Shekar, M., Wang, H., Park, J., Cui, W., et al. (2010). Ontology-based meta-analysis of global collections of high-throughput public data. PloS one 5.

Lai, Y. (2012). CCR5-targeted hematopoietic stem cell gene approaches for HIV disease: current progress and future prospects. Curr Stem Cell Res Ther 7, 310–317.

Lee, A.S., Tang, C., Rao, M.S., Weissman, I.L., and Wu, J.C. (2013a). Tumorigenicity as a clinical hurdle for pluripotent stem cell therapies. Nature Medicine 19, 998–1004.

Lee, M.O., Moon, S.H., Jeong, H.C., Yi, J.Y., Lee, T.H., Shim, S.H., Rhee, Y.H., Lee, S.H., Oh, S.J., Lee, M.Y., et al. (2013b). Inhibition of pluripotent stem cell-derived teratoma formation by small molecules. Proceedings of the National Academy of Sciences of the United States of America 110, E3281–3290.

Liao, S., Tammaro, M., and Yan, H. (2015). Enriching CRISPR-Cas9 targeted cells by co-targeting the HPRT gene. Nucleic acids research 43, e134.

Liu, J.C., Guan, X., Ryan, J.A., Rivera, A.G., Mock, C., Agrawal, V., Letai, A., Lerou, P.H., and Lahav, G. (2013). High mitochondrial priming sensitizes hESCs to DNA-damage-induced apoptosis. Cell stem cell 13, 483–491.

Mali, P., Yang, L., Esvelt, K.M., Aach, J., Guell, M., DiCarlo, J.E., Norville, J.E., and Church, G.M. (2013). RNA-guided human genome engineering via Cas9. Science 339, 823–826.

Merkle, F.T., and Eggan, K. (2013). Modeling human disease with pluripotent stem cells: from genome association to function. Cell stem cell 12, 656–668.

Mitzelfelt, K.A., McDermott-Roe, C., Grzybowski, M.N., Marquez, M., Kuo, C.T., Riedel, M., Lai, S., Choi, M.J., Kolander, K.D., Helbling, D., et al. (2017). Efficient Precision Genome Editing in iPSCs via Genetic Co-targeting with Selection. Stem cell reports 8, 491–499.

Musunuru, K. (2013). Genome editing of human pluripotent stem cells to generate human cellular disease models. Dis Model Mech 6, 896–904.

Nyquist, M.D., Corella, A., Burns, J., Coleman, I., Gao, S., Tharakan, R., Riggan, L., Cai, C., Corey, E., Nelson, P.S., et al. (2017). Exploiting AR-Regulated Drug Transport to Induce Sensitivity to the Survivin Inhibitor YM155. Mol Cancer Res 15, 521–531.

Radic-Sarikas, B., Halasz, M., Huber, K.V.M., Winter, G.E., Tsafou, K.P., Papamarkou, T., Brunak, S., Kolch, W., and Superti-Furga, G. (2017). Lapatinib potentiates cytotoxicity of YM155 in neuroblastoma via inhibition of the ABCB1 efflux transporter. Sci Rep 7, 3091.

Ramakrishna, S., Cho, S.W., Kim, S., Song, M., Gopalappa, R., Kim, J.S., and Kim, H. (2014). Surrogate reporter-based enrichment of cells containing RNA-guided Cas9 nuclease-induced mutations. Nature communications 5, 3378.

Rees, M.G., Seashore-Ludlow, B., Cheah, J.H., Adams, D.J., Price, E.V., Gill, S., Javaid, S., Coletti, M.E., Jones, V.L., Bodycombe, N.E., et al. (2016). Correlating chemical sensitivity and basal gene expression reveals mechanism of action. Nature chemical biology 12, 109–116.

Sadelain, M., Papapetrou, E.P., and Bushman, F.D. (2011). Safe harbours for the integration of new DNA in the human genome. Nat Rev Cancer 12, 51–58.

Seashore-Ludlow, B., Rees, M.G., Cheah, J.H., Cokol, M., Price, E.V., Coletti, M.E., Jones, V., Bodycombe, N.E., Soule, C.K., Gould, J., et al. (2015). Harnessing Connectivity in a Large-Scale Small-Molecule Sensitivity Dataset. Cancer discovery 5, 1210–1223.

Shi, Y., Inoue, H., Wu, J.C., and Yamanaka, S. (2017). Induced pluripotent stem cell technology: a decade of progress. Nat Rev Drug Discov 16, 115–130.

Sim, M.Y., Huynh, H., Go, M.L., and Yuen, J.S.P. (2017). Action of YM155 on clear cell renal cell carcinoma does not depend on survivin expression levels. PloS one 12, e0178168.

Steyer, B., Bu, Q., Cory, E., Jiang, K., Duong, S., Sinha, D., Steltzer, S., Gamm, D., Chang, Q., and Saha, K. (2018). Scarless Genome Editing of Human Pluripotent Stem Cells via Transient Puromycin Selection. Stem cell reports 10, 642–654.

Winter, G.E., Radic, B., Mayor-Ruiz, C., Blomen, V.A., Trefzer, C., Kandasamy, R.K., Huber, K.V.M., Gridling, M., Chen, D., Klampfl, T., et al. (2014). The solute carrier SLC35F2 enables YM155-mediated DNA damage toxicity. Nature chemical biology 10, 768–773.

Xie, Y., Wang, D., Lan, F., Wei, G., Ni, T., Chai, R., Liu, D., Hu, S., Li, M., Li, D., et al. (2017). An episomal vector-based CRISPR/Cas9 system for highly efficient gene knockout in human pluripotent stem cells. Sci Rep 7, 2320.

