## Supplemental Figure legends and Figures for "Scarless Enriched selection of Genome edited Human Pluripotent Stem Cells Using Induced Drug Resistance"

Running title: **Efficient scarless selection of edited hPSCs**

This PDF file includes:

Supplementary Materials and Methods

Supplementary Figures 1-5

Supplementary Table 1-2

### Supplementary Materials and Methods

#### Plasmid transfection and cell line establishment

For hESC transfections, 3 μg of Cas9 vector, 2 μg of sgRNA vector and 2 μg of siSLC35F2 were transfected through electroporation (NEPA-21). After 42 hours from transfection, cells were treated with 20nM of YM155 for 6 hours and added with media after DPBS washing for twice. Survived colonies were maintained and isolated through mechanical picking. Isolated colonies were analyzed by T7E1 assay and NGS.

#### Alkaline Phosphatase Staining

Alkaline phosphatase (AP) staining was performed with Alkaline Phosphatase Kit (86R1KT, Sigma Diagnostics™), following the supplier's instructions.

#### Fluorescence-based competitive proliferation assay

GFP-expressing hESC (EGFP-H9) and SLC35F2 KO hESC (SLC35F2 KO H9) are cultured together. Cells are detached with AccutaseTM (561527, BD Bioscience) and rinsed with DPBS three times before flow cytometry. GFP+ cells in the total population are measured using flow cytometry.

#### Immunocytochemistry

Cells were treated with 4% paraformaldehyde for fixation and 0.1% Triton X-100 in PBS for permeabilization. 5% normal serum albumin in PBS (blocking solution) is treated for 30 minutes for blocking. Primary antibody was diluted in blocking solution and applied to cells at 4°C, overnight. Primary antibody was washed off 3times with PBS. Secondary antibody and DAPI were diluted in blocking solution and applied to cells for 1 hour at room temperature, dark condition. Secondary antibody and DAPI were washed off 3times with PBS and mounted on slide glass using MOWIOL solution. Fluorescence microscopy (Olympus) was used for imaging samples.

**Targeted deep-sequencing analysis**

Genomic DNA from each samples was isolated using Wizard® genomic DNA purification kit (Promega). To assess gene editing, targeted deep sequencing near the DNA cleavage site was performed. Briefly, genomic DNA was prepared from the electroporated rotifer population, and the targeted regions (~ 300 bp) were amplified by adaptor primers with Phusion polymerase (New England Biolabs, Radnor, PA, USA). The detailed protocol was described in a previous study (Park et al., 2017). The resulting amplicons were subjected to paired-end read sequencing using MiniSeq (Illumina, San Diego, CA, USA). After MiniSeq, paired-end reads were joined by the Fastq-join utility, a part of the ea-utils program (https://code.google.com/archive/p/ea-utils/), using default values. Paired-end reads were then analyzed by comparing wild type and mutant sequences using CasAnalyzer (Park et al., 2017).

**RT-qPCR analysis**

Easy-BLUE$^{TM}$ RNA isolation kit (iNtRON Biotechnology) is used for total RNA extraction. PrimeScriptTM RT reagent kit (TaKaRa) is used to generate cDNA from RNA extracted previously. Quantitative real-time PCR analysis was performed with Light Cycler-480®II (Loche) and SYBR® Green PCR reagents (Life Technologies) are used for quantitative real-time PCR analysis, following the supplier's instructions.

**Immunoblotting**

Cells were lysed with Tissue lysis buffer or RIPA buffer with 10 uM sodium ortho-vanadate and 1 μM of protease inhibitor (Roche). Immuno-blotting was performed as described elsewhere.

**Supplementary Figure legends**

**Figure. S1** (A) Graphical summary for the induction of selective cell death mediated by YM155 in hPSCs mediated by SLC35F2 expression. (B) Annexin-V viability assay was performed in hPSCs followed by YM155 concentration. (C) Annexin-V cell death assay performed with SLC35F2 NC (Negative control) and KO #1 hPSCs after Bleomycin (BLM) treatment for 24 hrs. (D) Pluripotency-related proteins (LIN28A and SOX2) were immuno-stained with and images were taken through fluorescence microscopy. Insets are magnified images from each image (Scale bars = 200 μm). (E) Proliferation assay was performed with SLC35F2 NC and KO hPSCs through JuLi-Stage$^{TM}$. Representative images were taken by each time-point (0 hr, 24 hr, 48hr).

**Figure. S2** (A) Endogenous SLC35F2 gene expression profile among 15 normal cell lines was analyzed by NextBio portal (http://www.nextbio.com/b/nextbio.nb). (B) Annexin-V cell viability assay was performed with HEK293T cells followed by YM155 concentration. Assay was performed 24 hrs after YM155 treatment. (C) Representative images were taken after YM155 and Doxorubicin (DOXO) treated to SLC35F2 NC and KO 293T cells (scale bars = 200 μm) and (D) Annexin-V+ population was shown in histogram. (E) Intracellular amounts of YM155 in SLC35F2 NC and KO 293T cells were assayed and quantified. Red-dotted boxes indicate the specific peak for YM155. (F) Representative images were taken with 293T-surrogate cells after YM155 treatment to sgRNAs of CCR5 and SLC35F2 were treated. Red boxes are the cells only expressing mRFP while blue boxes are the cells with both expressing mRFP and GFP.

**Figure. S3** (A) T7E1 assay for CCR5 with enriched selection approach using YM155 in hESCs. (B) Mutation profiles of CCR5 target sequence were shown as a table based on deep-sequenced samples with Fig. S3A. The proportion of either insertion (Ins) or deletion (Del) mutation and overall indel ratio was calculated. (C, D) Independent experiment was performed as in Figs. S3A and S3B. T7E1 assay and deep-sequencing analysis was performed. (E) Graphs indicate the ratio of indel size of each population based on the deep-sequencing data from Figs S3C and S3D. (F) Sequence alignment of YES-selected clone targeting CCR5 as gene-of-interest. Control sequence indicates the target sequence of wild-type CCR5. Red box indicates the deletion mutation occurred in CCR5 targeted clone. (G) Alkaline phosphatase (AP) staining and gene expression profile of *SLC35F2* and pluripotency genes (*POU5F1*, *SOX2*, *NANOG*) was quantified (scale bars = 1 μm). (H) Representative images from of WT (wild type) and CCR5 targeted hPSCs after YM155 (10 nM or 20 nM) treatment for 24 hrs (scale bars = 200 μm).

**Movie. S1 Time-lapse video of WT and SLC35F2 KO hESCs after YM155 treatment**

**Movie. S2 Time-lapse video of 293T expressing the surrogate system**

**Movie. S3 Time-lapse video of 293T expressing the surrogate system after YM155 treatment**

**Table. S1 List of GSE studies for CTRP analysis**

**Table. S2 List of Gene Ontology terms significantly altered in SLC35F2 KO hESCs**

### References

Park, J., Lim, K., Kim, J.S., and Bae, S. (2017). Cas-analyzer: an online tool for assessing genome editing results using NGS data. Bioinformatics *33*, 286-288.

Figure. S1

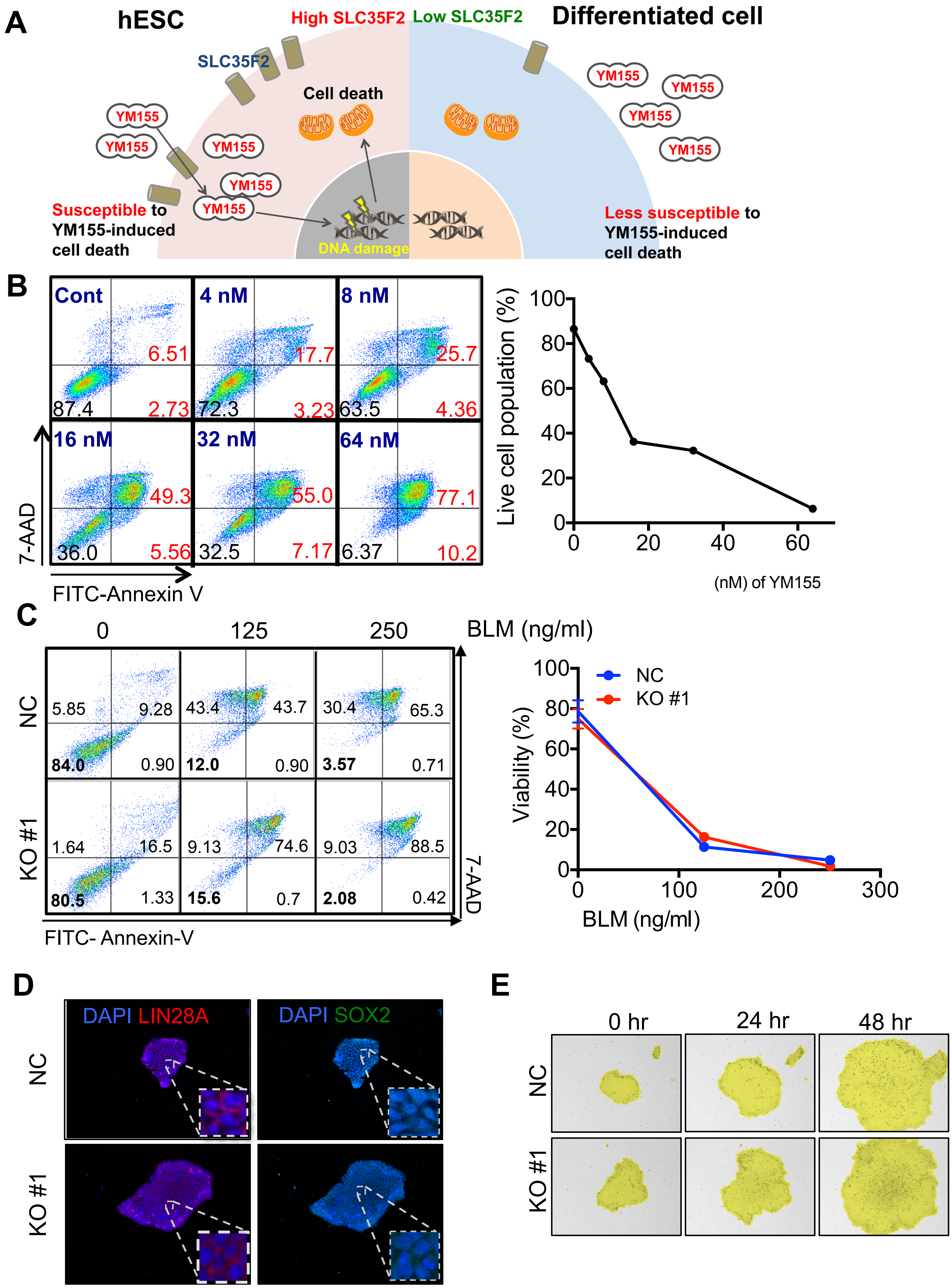

A
hESC
High SLC35F2
Low SLC35F2
Differentiated cell
SLC35F2
Cell death
YM155
DNA damage
Susceptible to YM155-induced cell death
Less susceptible to YM155-induced cell death
B
Cont
4 nM
8 nM
16 nM
32 nM
64 nM
6.51
87.4
2.73
17.7
72.3
3.23
25.7
63.5
4.36
49.3
36.0
5.56
55.0
32.5
7.17
77.1
6.37
10.2
7-AAD
FITC-Annexin V
Live cell population (%)
(nM) of YM155
C
0
125
250
BLM (ng/ml)
NC
KO #1
5.85
9.28
84.0
0.90
43.4
43.7
12.0
0.90
30.4
65.3
3.57
0.71
1.64
16.5
80.5
1.33
9.13
74.6
15.6
0.7
9.03
88.5
2.08
0.42
FITC- Annexin-V
7-AAD
Viability (%)
BLM (ng/ml)
D
DAPI LIN28A
DAPI SOX2
E
0 hr
24 hr
48 hr

Figure. S2

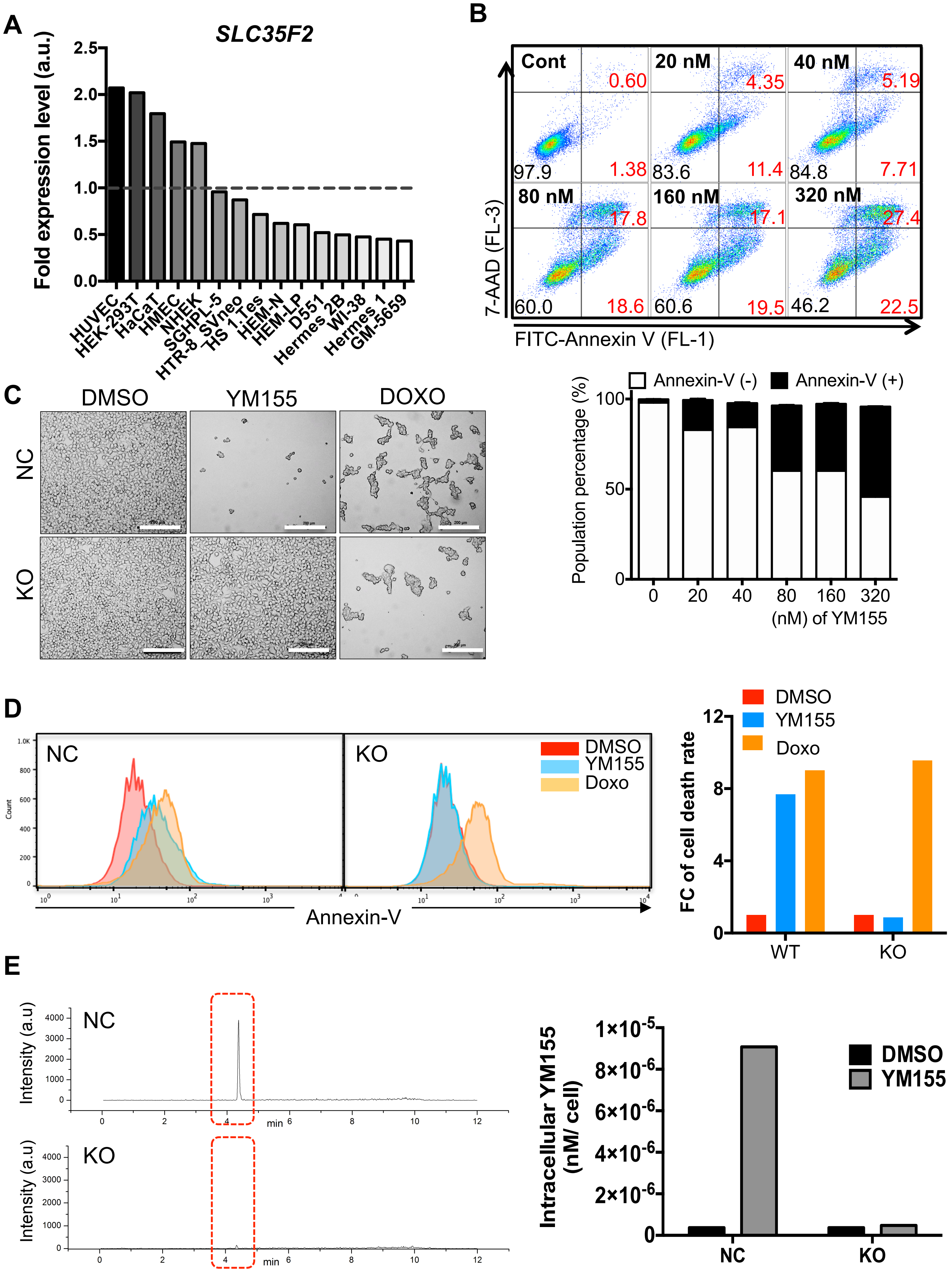
A
SLC35F2
Fold expression level (a.u.)
HUVEC
HEK-293T
HaCaT
HMEC
NHEK
SGHPL-5
HTR-8 SVneo
HS 1.Tes
HEM-N
HEM-LP
D551
Hermes 2B
WI-38
Hermes 1
GM-5659
B
Cont
20 nM
40 nM
80 nM
160 nM
320 nM
7-AAD (FL-3)
FITC-Annexin V (FL-1)
Annexin-V (-)
Annexin-V (+)
Population percentage (%)
(nM) of YM155
C
DMSO
YM155
DOXO
NC
KO
D
DMSO
YM155
Doxo
Count
Annexin-V
FC of cell death rate
WT
E
Intensity (a.u)
min
Intracellular YM155 (nM/ cell)

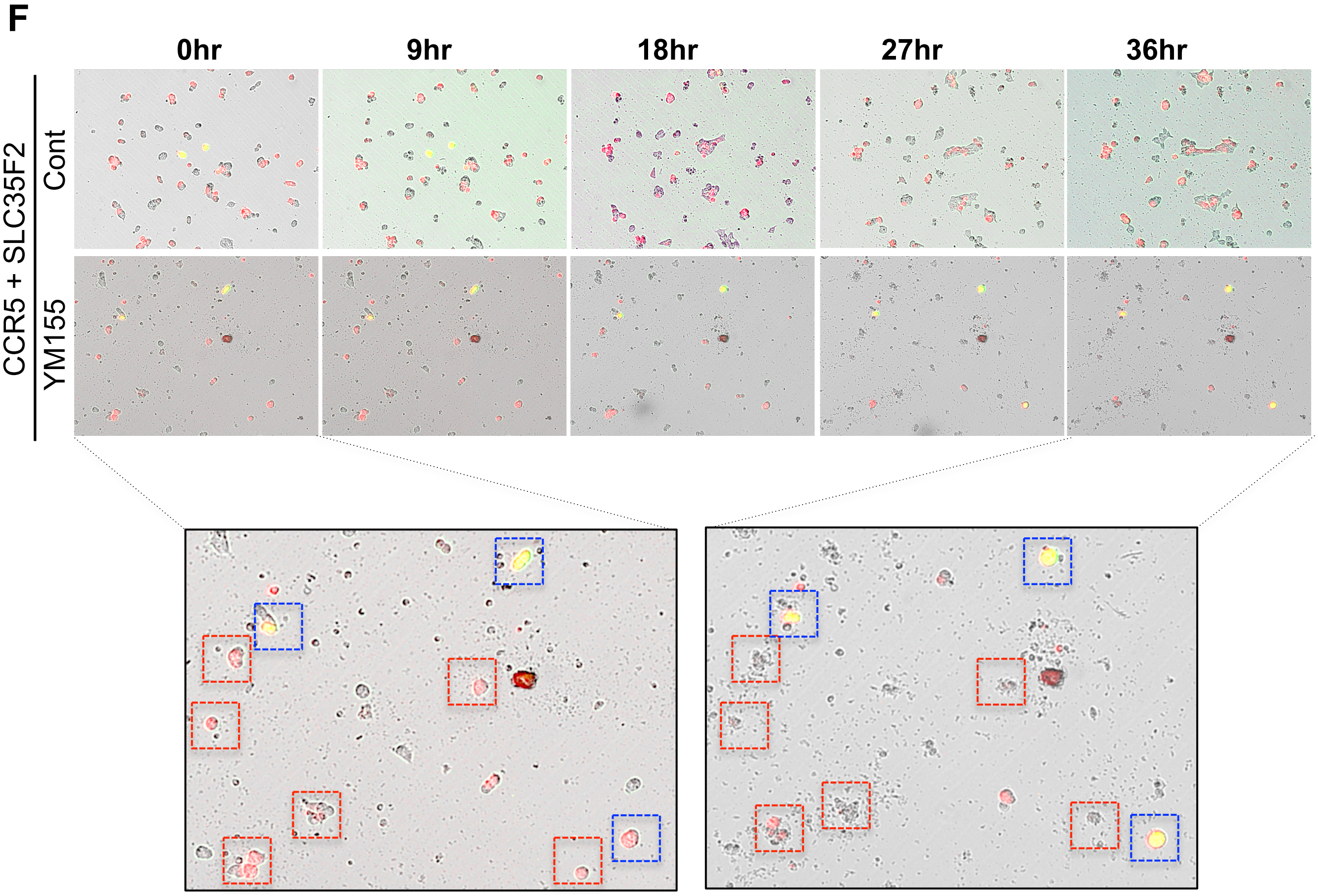

F
0hr
9hr
18hr
27hr
36hr
CCR5 + SLC35F2
Cont
YM155

### Figure. S3

**A**

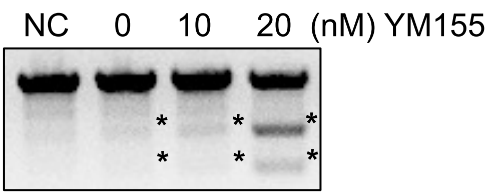


**B**

| Group | Reads | Ins | Del | Indel (Ratio) |
|---|---|---|---|---|
| Control | 64679 | 17 | 96 | 113 (0.2%) |
| DMSO | 64084 | 346 | 1168 | 1514 (2.4%) |
| 10 nM | 61998 | 503 | 2217 | 2720 (4.4%) |
| 20 nM | 33713 | 0 | 24328 | 24328 (72.2%) |

**C**

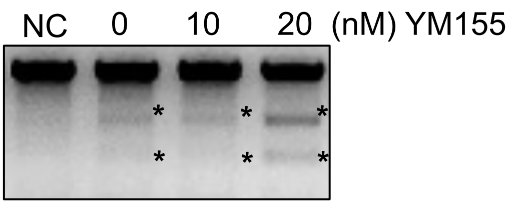


**D**

| Group | Reads | Ins | Del | Indel (Ratio) |
|---|---|---|---|---|
| Cont | 35336 | 2 | 81 | 83 (0.2%) |
| DMSO | 36605 | 225 | 861 | 1086 (3.0%) |
| 10 nM | 23668 | 229 | 511 | 740 (3.1%) |
| 20 nM | 37922 | 1196 | 5841 | 7037 (18.6%) |

**E**

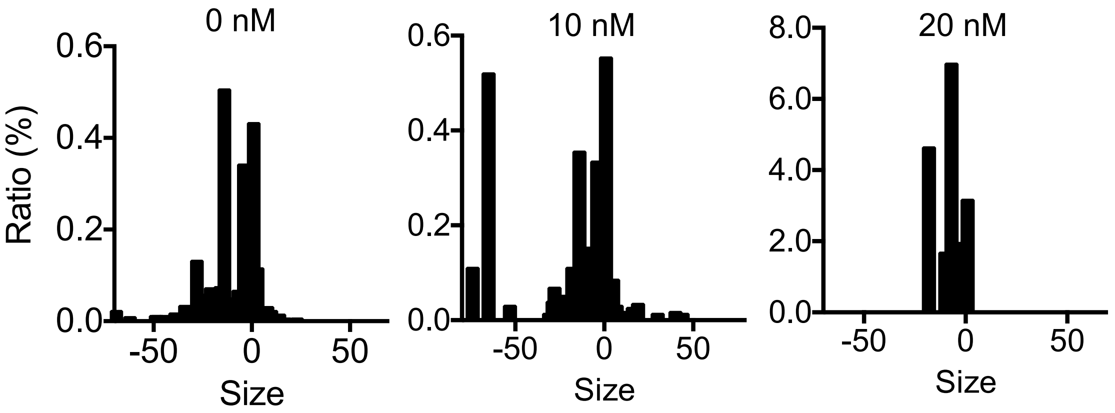


**F**

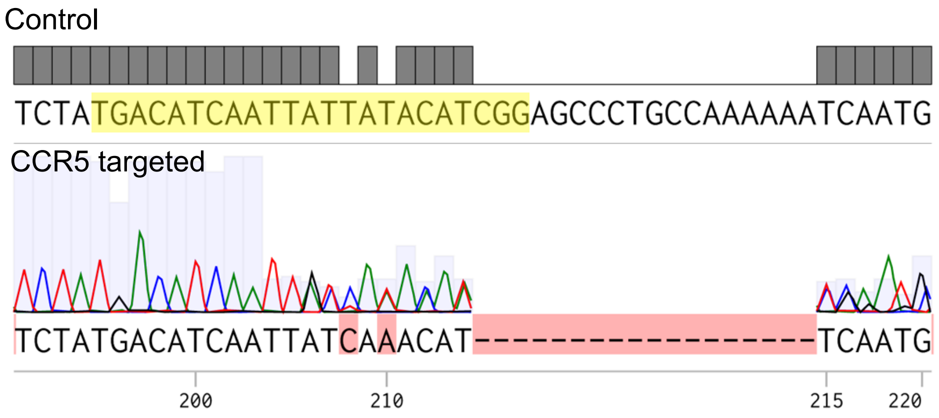

G

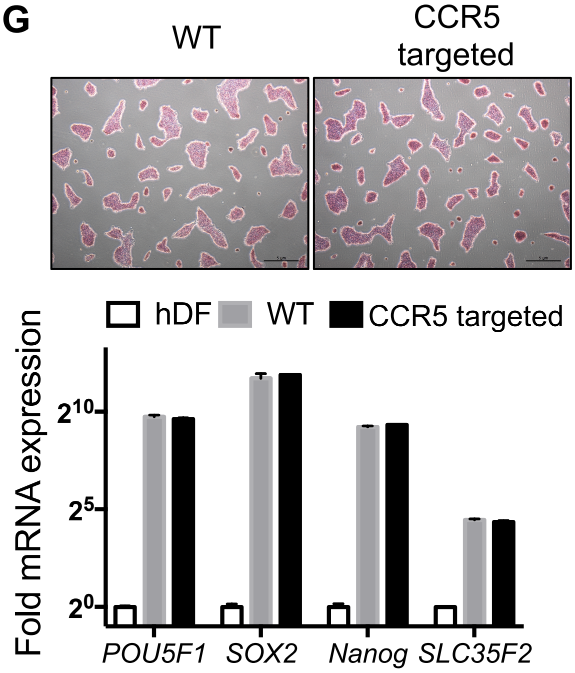
WT
CCR5 targeted
hDF
WT
CCR5 targeted
Fold mRNA expression
$2^{10}$
$2^{5}$
$2^{0}$
POU5F1
SOX2
Nanog
SLC35F2

H

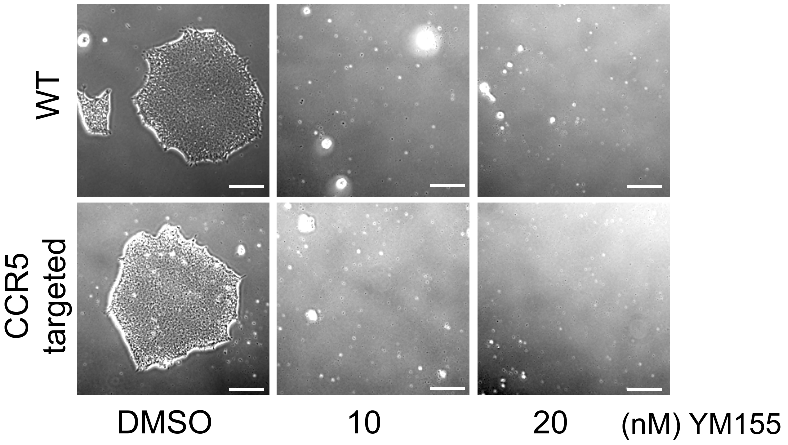
WT
CCR5 targeted
DMSO
10
20
(nM) YM155

| GSE # | Accession # | Cell type | Group | Samples |
|---|---|---|---|---|
| GSE28633 | GSM709608 | hESC | hPSC | 3 hESCs and 3 Neural cells |
| | GSM709609 | hESC | | |
| | GSM709610 | hESC | | |
| | GSM709617 | Neural cells | Differentaited | |
| | GSM709618 | Neural cells | | |
| | GSM709619 | Neural cells | | |
| GSE9709 | GSM245339 | hiPSC | hPSC | 3 hiPSCs and 2 hDF |
| | GSM248216 | hiPSC | | |
| | GSM257521 | hiPSC | | |
| | GSM245341 | hDF | Differentaited | |
| | GSM257254 | hDF | | |
| GSE23034 | GSM568484 | Hepatocyte- iPSCs | hPSC | 9 hiPSCs and 9 Differentiated (3 Hepatocyte, 3 Foreskin fibroblast, 3 Melanocyte) |
| | GSM568485 | Hepatocyte- iPSCs | | |
| | GSM568486 | Hepatocyte- iPSCs | | |
| | GSM568490 | Foreskin fribro-iPSCs | | |
| | GSM568491 | Foreskin fribro-iPSCs | | |
| | GSM568492 | Foreskin fribro-iPSCs | | |
| | GSM568496 | Melanocyte-iPSCs | | |
| | GSM568497 | Melanocyte-iPSCs | | |
| | GSM568498 | Melanocyte-iPSCs | | |
| | GSM568481 | Hepatocyte | Differentaited | |
| | GSM568482 | Hepatocyte | | |
| | GSM568483 | Hepatocyte | | |
| | GSM568487 | Foreskin Fibroblast | | |
| | GSM568488 | Foreskin Fibroblast | | |
| | GSM568489 | Foreskin Fibroblast | | |
| | GSM568493 | Melanocyte | | |
| | GSM568494 | Melanocyte | | |
| | GSM568495 | Melanocyte | | |
| GSE24487 | GSM603050 | BJ-iPSC | hPSC | 2 hESCs, 4 hiPSCs and 4 fibroblasts |
| | GSM603051 | BJ-iPSC | | |
| | GSM603052 | HGPS-iPSC | | |
| | GSM603053 | HGPS-iPSC | | |
| | GSM603054 | H9 | | |
| | GSM603055 | H9 | | |
| | GSM603015 | BJ fibroblast | Differentaited | |
| | GSM603043 | BJ fibroblast | | |
| | GSM603044 | HGPS fibroblast | | |
| | GSM603045 | HGPS fibroblast | | |
| GSE21073 | GSM525418 | iPSC-WT | hPSC | 3 hESCs, 6 hiPSCs and 2 hDFs |
| | GSM525419 | iPSC-WT | | |
| | GSM525420 | iPSC-WT | | |
| | GSM525421 | iPSC-WT | | |
| | GSM525422 | iPSC-WT | | |
| | GSM525423 | iPSC-WT | | |
| | GSM525424 | HUES6 | | |
| | GSM525425 | HUES6 | | |
| | GSM525426 | HUES6 | | |
| | GSM525415 | Fibroblast | Differentaited | |
| | GSM525416 | Fibroblast | | |
| | GSM525417 | Fibroblast | | |
| GSE25542 | GSM627785 | H9 | hPSC | 2 hESCs, 2hiPSCs and 9 Differentiated (8 Neurons, 1 fibroblast) |
| | GSM627767 | Fibroblast-iPSC | | |
| | GSM627791 | Fibroblast-iPSC | | |
| | GSM627792 | Fibroblast-iPSC | | |
| | GSM627782 | Neuron | Differentaited | |
| | GSM627783 | Neuron | | |
| | GSM627786 | Neuron | | |
| | GSM687788 | Neuron | | |
| | GSM627789 | Neuron | | |
| | GSM627793 | Fibroblast | | |
| | GSM627794 | Neuron | | |
| | GSM627795 | Neuron | | |
| | GSM627798 | Neuron | | |

| | Term | P-value | Z-score | Combined Score |
|---|---|---|---|---|
| Wiki Pathway | Zinc homeostasis_Homo sapiens_WP3529 | 2.2E-06 | -2.0 | 25.9 |
| | Copper homeostasis_Homo sapiens_WP3286 | 1.9E-03 | -2.1 | 13.1 |
| | Neural Crest Differentiation_Homo sapiens_WP2064 | 3.7E-03 | -1.9 | 10.5 |
| | Nuclear Receptors_Mus musculus_WP509 | 4.3E-03 | -1.9 | 10.3 |
| | Nuclear Receptors_Homo sapiens_WP170 | 4.9E-03 | -1.9 | 10.0 |
| KEGG pathway | Mineral absorption_Homo sapiens_hsa04978 | 4.7E-05 | -1.8 | 18.0 |
| | Neuroactive ligand-receptor interaction_Homo sapiens_hsa04080 | 7.5E-03 | -1.9 | 9.1 |
| | Morphine addiction_Homo sapiens_hsa05032 | 3.9E-02 | -1.8 | 5.8 |
| GO Molecular Function | sodium channel inhibitor activity (GO:0019871) | 0.033 | -3.36 | 11.49 |
| | gap junction channel activity (GO:0005243) | 0.013 | -2.62 | 11.47 |
| | olfactory receptor binding (GO:0031849) | 0.033 | -3.22 | 11.02 |
| | G-protein coupled peptide receptor activity (GO:0008528) | 0.002 | -1.70 | 10.91 |
| | wide pore channel activity (GO:0022829) | 0.016 | -2.57 | 10.67 |
| | neurotrophin binding (GO:0043121) | 0.064 | -3.82 | 10.46 |
| GO Biological Process | cellular response to zinc ion (GO:0071294) | 8.1E-07 | -2.07 | 29.1 |
| | response to zinc ion (GO:0010043) | 6.8E-07 | -2.04 | 29.0 |
| | zinc ion homeostasis (GO:0055069) | 4.9E-07 | -1.86 | 27.1 |
| | cellular response to cadmium ion (GO:0071276) | 2.7E-06 | -1.96 | 25.2 |
| | response to copper ion (GO:0046688) | 8.0E-06 | -2.10 | 24.7 |
